# Spatial transcriptomics reveals heterogeneous cell–cell interactions among brain regions in a cuprizone model consistent with multiple sclerosis lesions

**DOI:** 10.1101/2025.02.07.637125

**Authors:** Hui-Hsin Tsai, Sarbottam Piya, Jing Wang, Jing Zhu, Wenxing Hu, Andrew R. Gehrke, Shaolong Cao, Amanda J. Guise, Su Jing Chan, Mark Sheehan, Jenhwa Chu, Zhengyu Ouyang, Matthew Ryals, Michelle Lee, Wanli Wang, Edward Zhao, Patrick Cullen, Ravi Challa, Eric Marshall, Wanyong Zeng, Yea Jin Kaeser-Woo, Chris Ehrenfels, Luke Jandreski, Helen McLaughlin, Thomas M. Carlile, Jake Gagnon, Taylor L. Reynolds, Mingyao Li, Kejie Li, Baohong Zhang

## Abstract

The cuprizone (CPZ) model is widely used for modeling demyelination in multiple sclerosis (MS) and for testing potential remyelination therapies. We integrated single-cell and spatial transcriptomics (ST) to fine map the spatial cellular and molecular responses during de and remyelination. ST revealed global demyelination and neuroinflammation in the brain beyond the corpus callosum, with region-specific differences. We identified oligodendroglia and microglia as two major cell types with significant transcriptomic changes in the model. Ligand receptor pairing analyses predicted growth factor and phagocytic pathway enrichment during demyelination, which is consistent with changes in MS lesions. During remyelination, while mature oligodendrocytes nearly reversed their phenotype back to the control state, microglia remained associated with the demyelination phenotype. Finally, astrocytes in the CPZ model had the greatest preservation of disease-associated modules to MS lesions, while the MOL, OPC, and microglia showed moderate to low preservation, which overall suggested that the CPZ model had moderate translatability to chronically active MS lesions.

## Introduction

Multiple sclerosis (MS) is an inflammatory demyelinating disease of the central nervous system (CNS) affecting more than 2.8 million people worldwide ^1^. The pathological hallmarks of MS are gray and white matter demyelination, neurodegeneration, inflammation, and glial reactions ^2,3^. Some of the currently approved therapies have been shown to be very effective in reducing relapses associated with acute inflammation peripherally. However, despite well-controlled acute inflammation, disease progression can continue unabated, resulting in increased disability that lacks effective treatments ^4^. To better understand the mechanisms underlying the progressive biology of MS at the cellular and molecular levels, a recent genome-wide association study (GWAS) of the age-related MS severity score revealed that the DYSF–ZNF638 locus was associated with MS progression ^5^. Notably, the locus does not overlap with previously identified immunological susceptibility genes ^6^. The nearest gene to the disease severity locus, DYSF, encodes dysferlin, a protein specifically expressed by oligodendrocytes and neurons, underscoring CNS tissue involvement in MS disease progression. Parallel efforts have also been made to profile MS CNS tissue using emerging transcriptomic technologies to generate insights into MS disease pathogenesis ^7, 8, 9^.

Animal modeling is a critical part of understanding disease pathogenesis. The CPZ toxin model has been widely used to model demyelination and/or test potential new therapies for remyelination ^10^. CPZ is a copper chelator that inhibits mitochondrial functions. Mature oligodendrocytes (MOLs) demand high-energy metabolism ^11^ and are sensitive to CPZ-induced mitochondrial stress ^12^. The model has a stereotypical spatial-temporal demyelination-remyelination course, and the corpus callosum (CC) region shows a highly predictable demyelination pattern. During CPZ intoxication, demyelination and remyelination events may coexist at the same time as the lesion evolves until the peak of demyelination. This feature has been considered a strength of the model to mimic early-stage MS lesions where damage and repair can occur concurrently ^13, 14^. The CNS phenotypes in the CPZ model, such as demyelination, neurodegeneration, and gliosis, are traditionally characterized by histological staining, immunohistochemistry (IHC), and electron microscopy ^15, 16, 17^. While histopathological examination preserves the anatomical architecture, the disadvantage is that only a few selected markers can be examined, and the overall cellular and molecular response is not fully captured.

With the emerging transcriptomic data of single cells in the CPZ model ^18, 19^, we sought to further identify signatures associated with specific anatomical regions within the CNS by developing a methodology and workflow to analyze spatial transcriptomic data. We aimed to determine spatial expression profiles in the CPZ model, identify disease-associated cells in the CNS, and identify region-specific cell cell interactions that potentially underlie pathological processes. MS lesions are heterogeneous in nature, and the location for lesion formation is unpredictable ^20^. The methodology developed in this study can be applied to human MS samples in the future to map expression patterns and changes in the lesion environment.

Here, we resolved spatial gene expression in the CPZ model by employing spatial transcriptomic (ST) technology ^21, 22^ and integrating the ST data with single-nucleus RNA-seq (snRNA-seq) references from the same study. We found that demyelination and neuroinflammation were prominent and widespread in the CPZ brain, and interestingly, some biological pathways showed discordant directionality in different brain regions. Ligand receptor (LR) pairing analyses revealed enrichment of phagocytic and growth factor pathways in the CPZ group from both the snRNA-seq and ST data. Finally, we compared the snRNA-seq data from the CPZ model with a human MS dataset ^7^ and found moderate levels of preservation of MS-associated disease modules in the CPZ model.

## Results

We established three experimental groups and collected tissues from similar brain regions (approximately bregma -1.5 mm) for different sequencing platforms, that is, snRNA-seq, bulk RNA-seq, and 10X Visium spatial gene expression profiling (Figure 1A, Methods). We fed 8- to 10-week-old mice 0.3% CPZ for 4 weeks to induce peak demyelination. The recovery group (RCV) received 4 weeks of CPZ followed by 2 weeks of normal chow to allow spontaneous remyelination. The control group received normal chow for 6 weeks.

**Figure 1.**
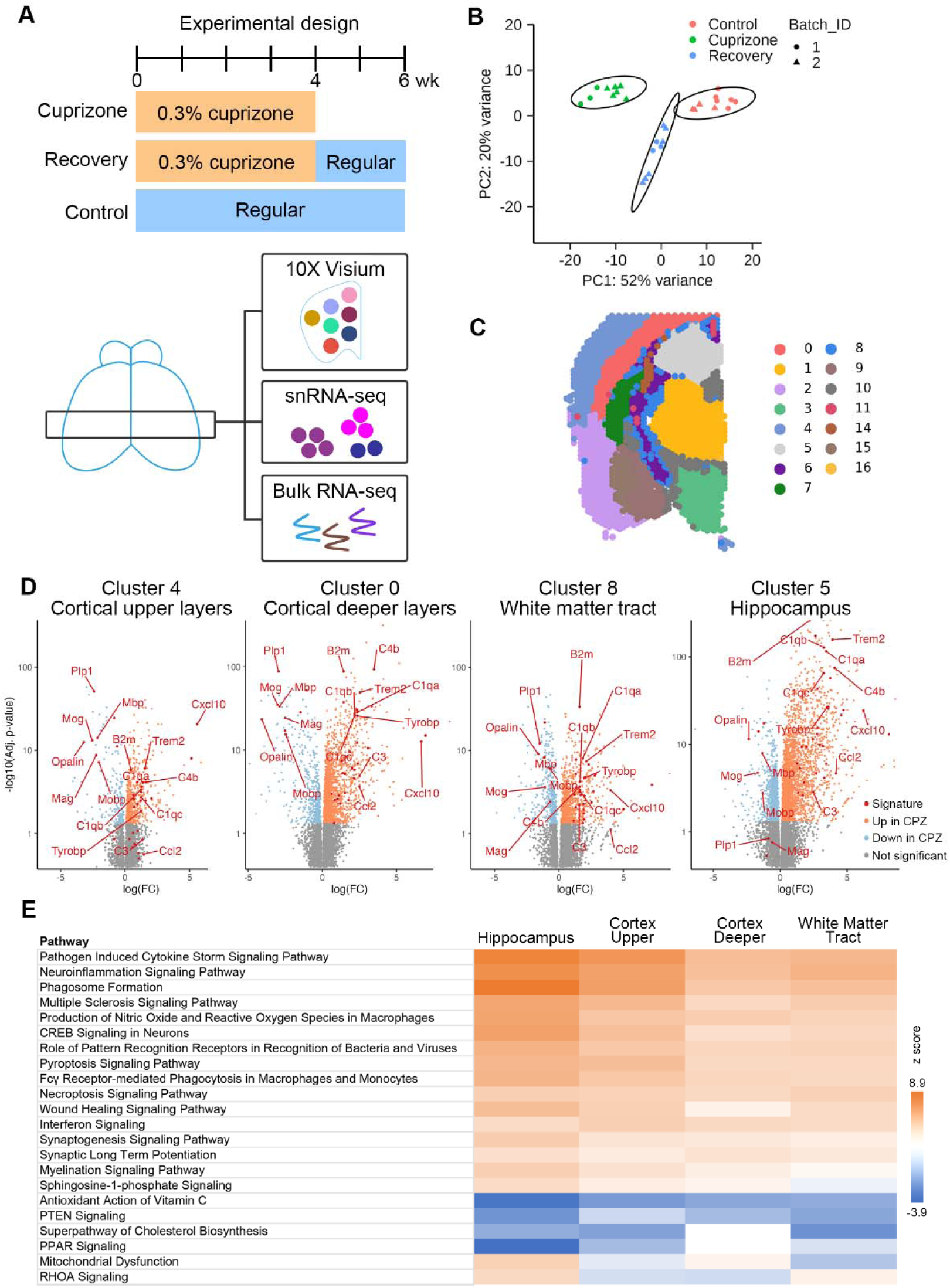
Widespread demyelination and neuroinflammation occur along the dorsal-ventral axis of the CPZ brain. (A) CPZ model experimental design. Three experimental groups were collected in this study, and brain tissue was collected for 3 sequencing platforms. (B) Spatial transcriptomic data from three experimental groups were distinctly separated on the PCA plot after pseudobulking. PCA plot of spot-level pseudobulk across biological replicates from the ST data, with each point representing a biological replicate. The point color represents the experimental conditions, and the point shape represents the batch. (C) The 17 clusters of ST spots identified by the SpaGCN algorithm strongly resembled the reference anatomical annotations (Allen Brain Atlas mouse reference atlas: https://mouse.brain-map.org/). (D) Volcano plots of differentially expressed genes (DEGs) between CPZ and control conditions in cluster 4 (cortical upper layers), cluster 0 (cortical deeper layers), cluster 8 (white matter tract) and cluster 5 (hippocampus) are labeled with a list of curated MS signature genes. The negative log adjusted p values (y-axis) are presented on a log scale. (E) The majority of pathways showed congruous directionality among clusters, while some showed incongruous directionality. DEGs in each cluster were analyzed by IPA core analysis for pathway discovery and subsequently subjected to comparison analysis.

### ST revealed global demyelination and region-specific transcriptomic changes induced by CPZ

We first compared the Visium spatial maps of the Mbp, Cd68, and Gfap genes at the spot level via IHC staining and showed that the expression patterns within the sections were comparable between the Visium and IHC data (Figure S1). These genes were chosen based on their well-established expression patterns during de and remyelination in the CPZ model. The ST dataset of all sections contained a total of 57,217 spots (average = 2043), with an average of 75,141 reads per spot and a median of 4,157 genes per spot. To look at the overall structure of the ST dataset, we performed unsupervised spot-level pseudobulk principal component analysis (PCA) by aggregating spot counts for each gene across all spots within each section to form a gene-by-section count matrix. Our results showed that the three experimental groups were distinctly separated along PC1 (Figure 1B). We also observed concordance of cell abundance estimates in the same region between animals after ST cell type deconvolution (Figure S2), which further suggested that the separation on PCA was driven by experimental groups.

We performed unbiased clustering of the ST spots using the SpaGCN package ^23^ and identified 17 spatial clusters closely resembling the reference anatomical regions (http://www.brain-map.org) (Figure 1C). We identified 484, 1495, 1086 and 2762 differentially expressed genes (DEGs) between CPZ and control conditions in Cluster 4 (occupying cortical upper layers, roughly layers I-III), Cluster 0 (occupying cortical deeper layers, roughly layers IV-VI), Cluster 8 (occupying white matter tract), and Cluster 5 (occupying hippocampus), respectively. Cluster 5 (hippocampus) had the greatest number of DEGs and highest fold changes among the 4 clusters (Figure 1D). We curated a list of genes that represent the gene signature of MS ^7, 9^ and found that proinflammatory genes (B2m, Cxcl10, C4b, and Trem2) were upregulated and that myelin-related genes (Mbp and Mog) were downregulated in all 4 clusters (Figure 1D). While the CC has been viewed as the major anatomical location for evaluating demyelination and remyelination in CPZ model ^24, 25^, our analysis showed that inflammatory and demyelination signatures were prominent and widespread in various brain regions. By comparing clusters in close anatomical proximity, such as Clusters 4 and 0, we also found that more genes in the MS gene signatures reached statistical significance in Cluster 0, suggesting a more prominent inflammatory state in deeper cortical layers. This observation parallels a previous finding that the upper cortical layers remyelinated more robustly than did the deeper layers in the CPZ model, and the difference in the repair capacity was attributed to the hypothesized increase in inflammatory activity, myelin debris, and gliosis in the deeper layers ^26^.

Further analysis of biological pathways revealed that neuroinflammation and cell death pathways were prominently upregulated in all 4 clusters/regions. PTEN signaling and the superpathway of cholesterol biosynthesis were downregulated in all 4 clusters, and they have both been linked to myelin integrity and demyelination ^27, 28^ (Figure 1E). Interestingly, both mitochondrial dysfunction and RhoA signaling, which are implicated in MS pathology, were up- and downregulated depending on the cluster or anatomical region.

### Oligodendroglia and microglia were the major cell types associated with the CPZ model

To resolve the cell types at each ST spot, we collected samples from dorsal brain regions, including the cortex, CC, and hippocampus, to perform snRNA-seq. We obtained a total of 294,447 nuclei with a mean of 3,392 detected genes per nucleus after filtering out low-quality cells and potential doublets and removing ambient RNAs. By transferring cell labels from publicly available datasets, we identified a total of 32 annotated cell types (Figure 2A) ^29, 30, 31^.

**Figure 2.**
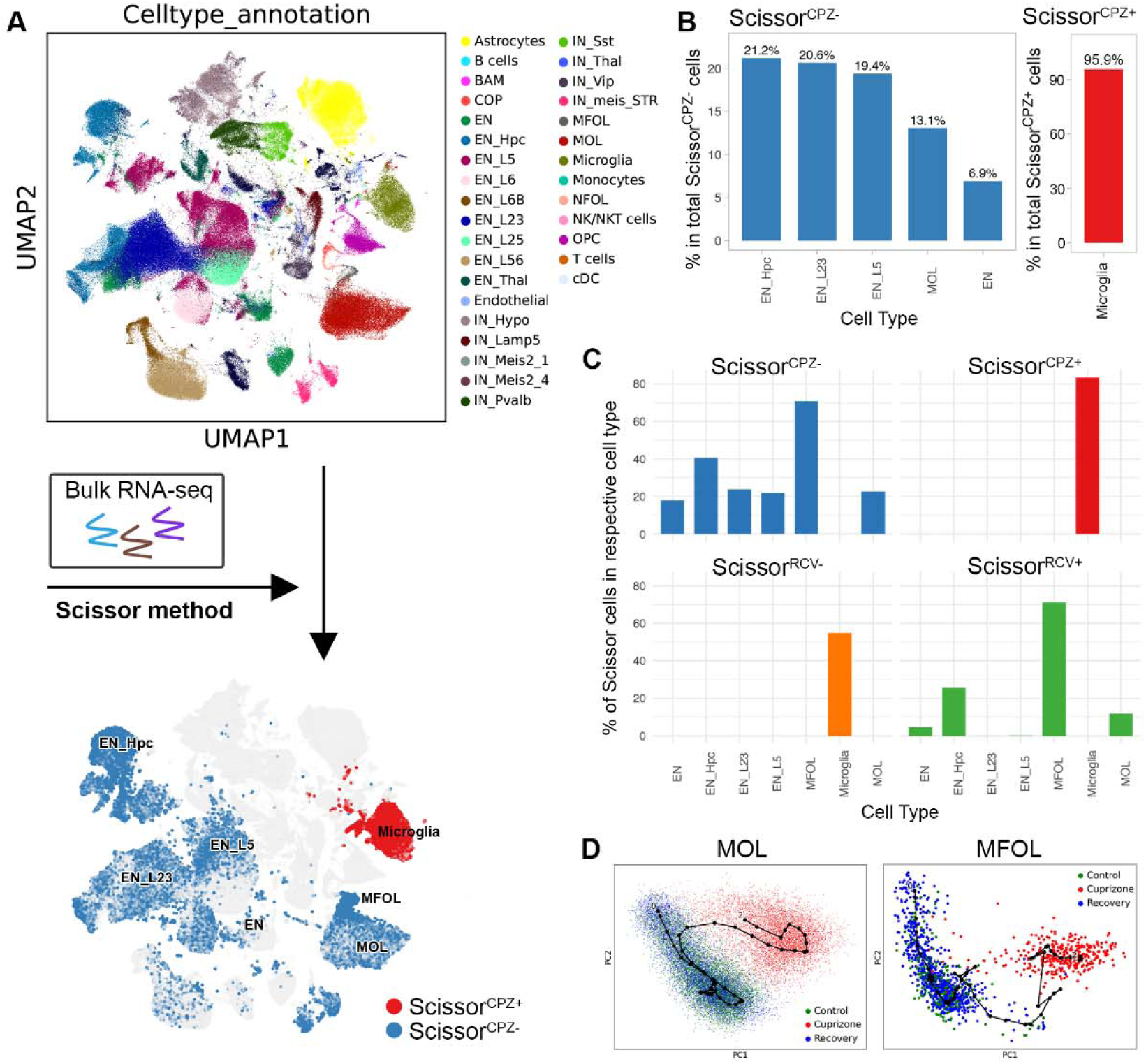
Identification of CPZ-associated cell types by integrating bulk and snRNA-Seq data with Scissor. (A) Single-cell UMAP projection of the mouse brain snRNA-Seq data colored by cell annotation. By integrating bulk RNA-seq data, blue Scissor^CPZ-^ and red Scissor^CPZ+^ were identified. (B) Bar plots demonstrating a breakdown of the cell type compositions within the total of Scissor^CPZ-^ or Scissor^CPZ+^ cells. (C) Bar plots showing a detailed breakdown of the percentage of Scissor cells in each cell type. (D) Pseudotime analysis of snRNA-Seq data for MOL and MFOL showed that their CPZ phenotypes were distinct from those of the control and RCV phenotypes.

In addition, we collected matched samples for bulk RNA-seq. We integrated both bulk and single-cell data using the Scissor package to unbiasedly identify cell populations associated with experimental phenotypes. This method does not require unsupervised clustering of single-cell data, which avoids subjective decisions regarding cell cluster numbers or clustering resolution ^32^. We found that 2.99% of cells (8,802 cells among 294,447 nuclei) were positively associated with the CPZ phenotype (Scissor^CPZ+^ cells), 14.9% of cells (43,891 cells) were negatively associated with the CPZ phenotype (Scissor^CPZ-^ cells), and the remaining cells were considered background cells (not significantly associated with the CPZ phenotype) (Figure 2B and 2C). Among Scissor^CPZ+^ cells, 95.9% were microglia and within the microglial population, 83.3% were classified as Scissor^CPZ+^. Therefore, a high percentage of microglial cells contributed to Scissor^CPZ+^ cells. Among Scissor^CPZ-^ cells, 13.1% were MOL, 2.14% were MFOL (myelin-forming oligodendrocytes), 21.2% were EN_Hpc, 20.6% were EN_L23, 19.4% were EN_L5 and 6.9% were EN. Furthermore, 70.8% of MFOL, 40.6% of EN_Hpc, 22% of MOL, 23.8% of EN_L23, 21.9% of EN_L5 and 18% of EN were classified as Scissor^CPZ-^ within their respective cell type populations (Figure 2C). In the recovery group, 4.13% of the cells (12,182 cells) were positively associated with the RCV phenotype (Scissor^RCV+^ cells), and 1.95% of the cells (5,729 cells) were negatively associated with the RCV phenotype (Scissor^RCV-^ cells). The Scissor^RCV+^ cells comprised 25% MOL, 48% EN_Hpc, and 7.8% MFOL, and the Scissor^RCV-^ cells comprised 96.6% microglia. MFOL had the highest percentage being classified as Scissor^RCV+^ and microglia had the highest percentage being classified as Scissor^RCV-^ (Figure 2C). Among the Scissor cell types, microglia had the greatest number of DEGs, and neurons generally had a lower number of DEGs (Figure S3).

The results from Scissor suggested that oligodendroglia and microglia were the major cell types associated with the CPZ model. Moreover, in the RCV group, microglia exhibited a sustained demyelination phenotype, while oligodendroglia exhibited a phenotype closer to that of the control group. The similarity of oligodendroglia between the RCV and control phenotypes was further supported by pseudotime analysis. That is, the CPZ phenotype was distant from the RCV and control phenotypes, while the recovery and control phenotypes were indistinguishable from each other (Figure 2D).

### Identification of oligodendroglia and microglial subclusters associated with CPZ phenotypes

We focused on oligodendroglia and microglia based on Scissor findings and identified 12 subclusters from oligodendroglia, including subclusters 0, 1, 3, 4, 6, 9, 10, and 11 from MOLs; subclusters 3, 4, and 6 from MFOLs; subclusters 5 and 7 from NFOLs (newly formed oligodendrocytes); subcluster 5 from COPs (differentiation-committed oligodendrocyte precursors); and subclusters 2 and 8 from OPCs (oligodendrocyte precursors) (Figure 3A). The proportion of cells in subcluster 3 increased significantly in the CPZ condition (CPZ vs. control, p value = 6.223e-10) and decreased significantly in the RCV condition (RCV vs. CPZ, p value = 5.861e-10) (Figure 3B and 3C). This result supported that subcluster 3 was exclusively associated with the CPZ condition and expressed the marker genes Arap2, Dock10, Tenm4, Pex5l and Dock1 (Figure 3D). On the other hand, subclusters 0, 1, and 4 showed a marked reduction in proportion in the CPZ treatment (CPZ vs. control, p value = 0.006, 0.0004, and 0.0475, respectively). The proportions of these 3 subclusters increased in the RCV condition, suggesting a phenotype reversal, i.e., remyelination (RCV vs. CPZ, p value = 0.03, 0.001, and 0.002, respectively) (Figure 3C).

**Figure 3.**
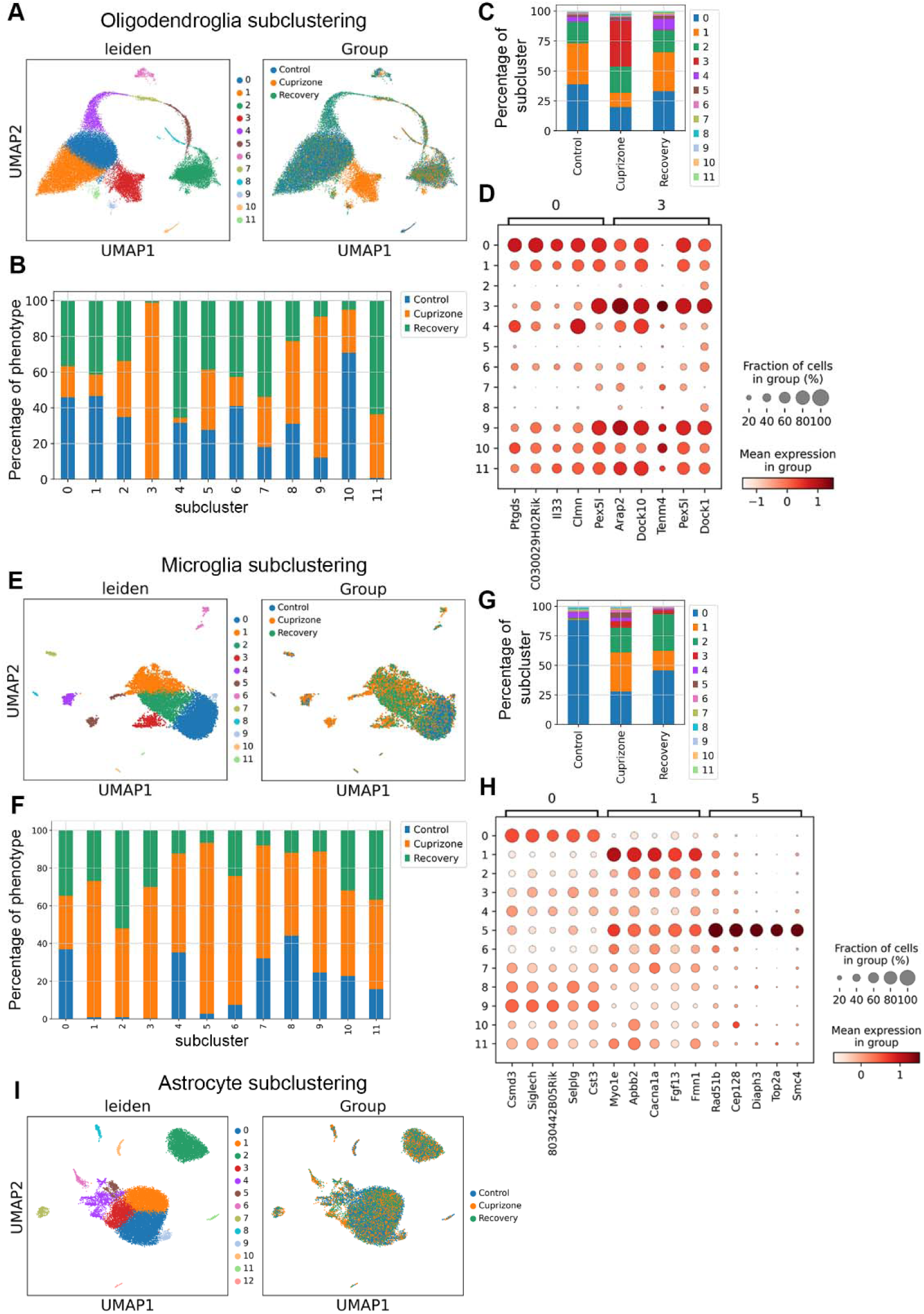
Identification of CPZ-associated oligodendroglia and microglial subclusters. (A) The UMAP of oligodendroglia is colored according to subclusters and are separated by phenotype. (B) Stacked bar plot of oligodendroglial subclusters according to treatment phenotype. (C) Stacked bar plot of treatment phenotypes according to the percentages of oligodendroglial subclusters. (D) Dot plot of the top 5 marker genes specific to subcluster 0 and subcluster 3 (CPZ-associated subclusters) of oligodendroglia. (E) The UMAP of microglia is colored according to subcluster and are separated by phenotype. (F) Stacked bar plot of microglial subclusters according to treatment phenotype. (G) Stacked bar plot of treatment phenotypes colored according to the percentages of microglial subclusters. (H) Dot plot showing the top 5 marker genes specific to subcluster 0 (homeostatic subcluster) and subclusters 1 and 5 (CPZ-associated subclusters) of microglia. (I) UMAP of astrocytes colored according to subcluster did not show separation according to phenotype.

We also identified 12 subclusters from microglia (Figure 3E to 3H). More than 88% of the control microglia were assigned to subcluster 0, which expressed homeostatic genes, such as Tmem119, P2ry12, Tgfbr1, and Siglech. Microglia expanded substantially during demyelination, and the proportions of microglia in the control, CPZ and RCV groups were 19%, 46.5% and 34.5%, respectively. CPZ-associated subcluster 1 showed high expression of the ApoE, Axl, Cd9 and Lpl genes, which have been implicated as neurodegenerative disease-associated genes ^33^. The proportion of subclusters 1 and 5 exhibited a significant increase during demyelination (CPZ vs. control, p value = 1.567e-08 and 0.029, respectively), followed by a significant decrease during remyelination (RCV vs. CPZ, p value = 0.009 and 0.035, respectively) (Figure 3G). Unlike those in oligodendroglia, the changes in the proportions of microglial subclusters from the CPZ to the RCV are far from normalization compared to those in control conditions. In contrast, we did not find any subclusters of astrocytes associated with the CPZ phenotype (Figure 3I).

### CellChat identified phagocytosis, growth factor, and chemokine pathways as novel cell-cell interactions emerging in CPZ condition from snRNA-seq

We analyzed the snRNA-seq dataset by CellChat ^34^ to uncover novel enriched ligand-receptor (LR) pairs, focusing on predicting significantly altered interactions to CPZ and control contrast.

We identified enhanced phagocytosis, growth factor, and chemokine pathways in microglia, either as receiver or emitter cells under the CPZ condition (Figure 4A and 4B). When microglia were analyzed as receiver cells, they paired with multiple emitter cell types with significant novel LR pairs, such as GAS (Gas6-Mertk), NRG (Nrg1-(Itgav+Itgb3)), CX3C (Cx3cl1-Cx3cr1), VEGF (Vegfa-Vegfr1), ANGPT (Angpt2-(Itga5+Itgb1)), and ANGPTL (Angptl2-(Itga5+Itgb1)) (Figure 4A). The Gas6-Mertk pathway is considered an anti-inflammatory pathway and has been shown to be critical for modulating microglial phagocytic activity to remove myelin debris, therefore promoting remyelination in the CPZ model ^35^. Interestingly, pathways involved in blood vessel remodeling, such as the VEGF and ANGPT pathways, were predicted to be present in microglia ^36, 37^. Increasing evidence has suggested that interactions between microglia and blood vessels could play roles in diseases ^38, 39^. Understanding the functional significance of the VEGF and ANGPT pathways in microglia and how they contribute to demyelination and/or remyelination processes or even blood vessel homeostasis may provide new insights into MS disease pathology ^37^. Other novel LR pairs in the CPZ condition included FGF (Fgf1-Fgfr2, astrocytes to microglia), GDN (Gdf15-Tgfbr2, MOL to microglia), and BMP (Bmp7-(Acvr1+Bmpr2), OPC to microglia) pairs. Growth differentiation factor 15 (Gdf15) is a neurotrophic factor of the TGFβ superfamily, and elevated levels of Gdf15 in serum are associated with MS disease stability and are presumed to be anti-inflammatory ^40^. Gdf15 was recently identified as a marker of demyelination-associated oligodendrocytes and was specific to the CPZ model but not the 5XFAD model of Alzheimer’s disease ^19^. Here, we predicted that Gdf15 expression in the MOL may exert its function through Tgfbr2 on microglia.

**Figure 4.**
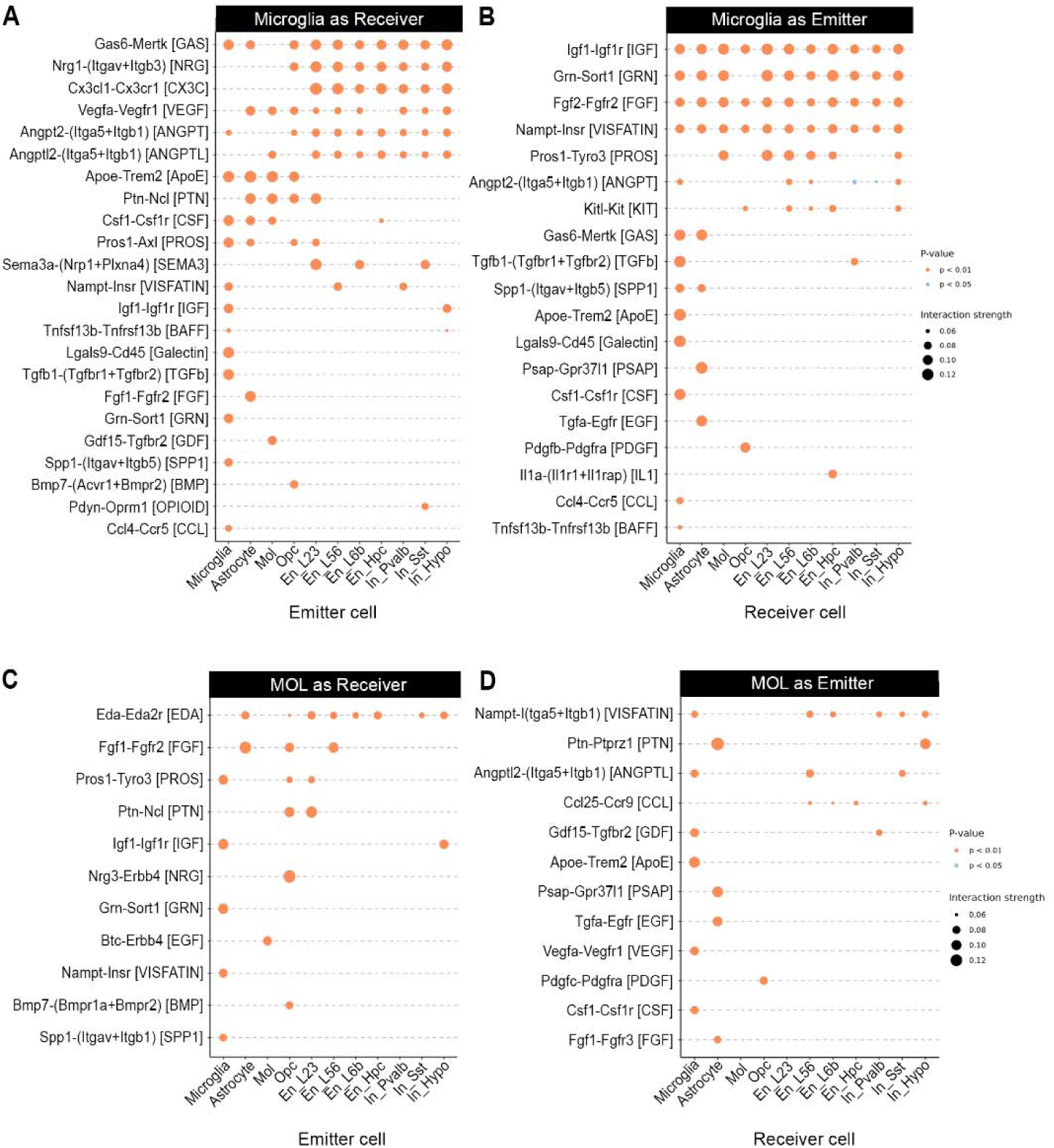
Novel LR pairs under the CPZ condition predicted from the single-cell dataset by CellChat. (A) CellChat analysis of the snRNA-seq data predicted novel CPZ LR pairs with microglia as the receiver. The associated CellChat pathway information is listed in brackets beside the LR name along the vertical axis and sender cell types along the horizontal axis. The point size represents the CellChat interaction strength. The point color indicates the significance cutoff. (B) Novel LR pairs with microglia as sender cells. (C) Novel LR pairs with MOL as a receiver cell. (D) Novel LR pairs with MOL as a sender cell.

When microglia were analyzed as emitter cells, interactions between IGF (Igf1-Igf1r), GRN (Grn-Sort1), FGF (Fgf2-Fgfr2), VISFATIN (Nampt-Insr), and PROS (Pros1-Tyro3) were predicted in multiple receiver cell types (Figure 4B). PSAP (Psap-Gpr37l1, microglia to astrocytes), EGF (Tgfa-Egfr, microglia to astrocytes), PDGF (Pdgfb-Pdgfra, microglia to OPC), and IL1 (Il1a-(Il1r1+Il1rap), microglia to EN-Hpc) pathway interactions were predicted between microglia and selective cell types. Finally, TGFb (Tgfb1-(Tgfbr1+Tgfbr2)), ApoE (ApoE-Trem2) and Csf1-Csf1r (CSF) were identified as pathway interactions between microglia. Among these pathways, most have been demonstrated to be neuroprotective, while the Csf1-Csf1r interaction is considered proinflammatory. For example, the IGF pathway could play a role in decreasing the severity of neuroinflammation ^41^, and Gpr37l1 expression in astrocytes modulates neuronal NMDA receptor activation during ischemia to exert protective effects ^42^. So far, LR analysis by CellChat revealed that microglia interacting pairs may provide protective functions in addition to proinflammatory functions under CPZ conditions.

Most of the LR pairs identified from MOL were involved in growth factor signaling pathways (Figure 4C and 4D), such as the FGF, EGF, IGF, and VEGF pathways, regardless of whether the MOL was a receiver or emitter cell type. Interestingly, our analysis also revealed MOL and OPC interactions under CPZ conditions. When MOLs were analyzed as receiver cells, they received FGF, PTN and NRG signaling from OPCs, whereas OPCs received PDGF signaling from MOLs (Pdgfc-Pdgfra, OPC to MOL) when the MOLs were analyzed as emitter cells. To our knowledge, the cell cell interaction between MOLs and OPCs has not been well established, especially under demyelination conditions, which could be important for understanding the repair capacity of oligodendrocytes in diseases.

### ST mapped CPZ-associated cell subclusters to CC

Since the 10X Visium Spatial Gene Expression platform captures multiple cells per spot, we used the snRNA-seq data as a reference to deconvolute cell types at each spot by cell2location (Figure 5A) ^29^. As expected, EN-Hpc neurons were mapped to the hippocampal region after deconvolution (Figure 5A). To take advantage of ST to resolve gene expression in each cell type spatially, we manually annotated regions of interest (the CC, cortex, and hippocampus) in each Visium section for subsequent analyses based on available single-cell data from those regions (Figure 5A).

**Figure 5.**
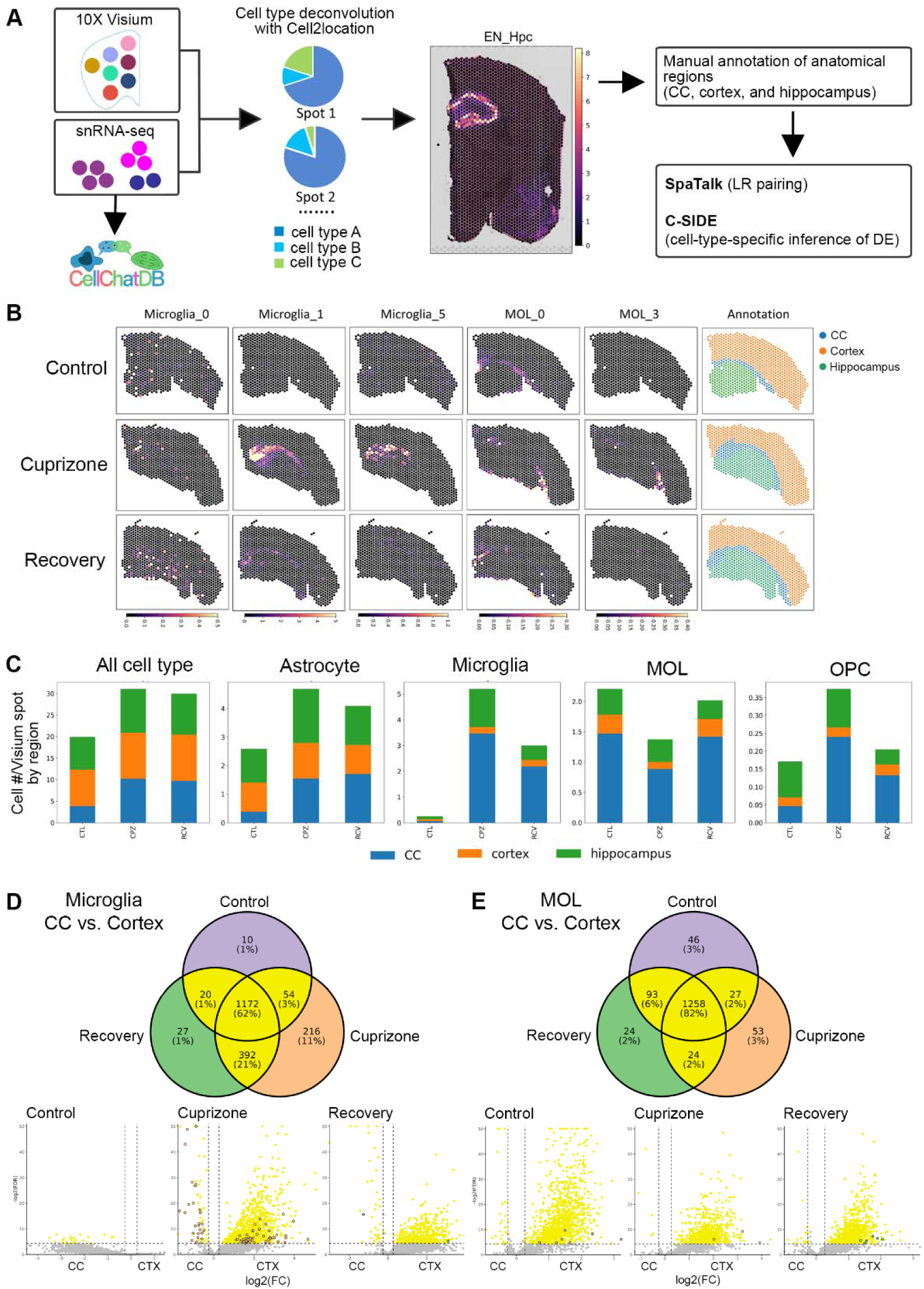
Deconvoluted ST data mapped CPZ-associated cell subclusters and demonstrated cell type abundance and expression heterogeneity across brain regions. (A) The workflow illustrating the steps involved in the ST data analysis. (B) Spatial plot showing the abundance of CPZ-associated subclusters of microglia and oligodendrocytes using cell2location. (C) Stacked bar plot representing the abundance of different cell types across various treatments in different brain regions. (D) Differential expression of microglial genes in CC compared to the cortex across different treatments represented by genes detected in each condition in the Venn diagram (upper panel) and DEGS volcano plots (lower panel), with yellow genes being DEGs shared in at least 2 conditions and purple, orange, and green points representing DEGs unique to CTL, CPZ, and RCV respectively, and (E) Differential expression of MOL genes in the CC compared to the cortex across different treatments represented by Venn diagram (upper panel) and volcano plots (lower panel) as described in D.

CPZ-associated microglial subclusters 1 and 5 were mapped mostly to the medial CC on the representative CPZ section, and these subclusters were almost nonexistent on the control or RCV sections (Figure 5B). This finding is consistent with the IHC data showing that microglial activation is most prominent in the CC under CPZ conditions (Figure S1). In contrast, homeostatic microglia subcluster 0 did not concentrate in any particular region. CPZ-associated oligodendroglia subcluster 3 was mapped mostly onto the lateral CC on the representative CPZ section and was almost nonexistent on the control or RCV sections. Oligodendroglia subcluster 0 was mapped to the entire CC region in the control section and to the medial CC region in the RCV section. Interestingly, MFOLs from the oligodendrocyte lineage mostly mapped to the RCV section (Figure S4A), suggesting an active remyelination process.

We next investigated the abundance of each cell type by region under each condition. In general, there was a substantial increase in the number of cells per Visium spot in the CC (Figure 5C). Microglia exhibited a sharp increase in cell abundance in the CC and hippocampus under CPZ conditions, and the abundance remained high under RCV conditions, suggesting incomplete reversal of the phenotype to that of the control. On the other hand, the MOL abundance in the CC, cortex, and hippocampus was reduced in the CPZ group but recovered to the control level in the RCV group, suggesting a robust remyelination process after CPZ withdrawal. The OPC number in the CC continued to remain high under RCV conditions.

Finally, we investigated region-specific cellular responses of microglia and the MOL. Microglial populations in the cortex and CC had their own gene signatures under control conditions. While microglia in the CC expressed more genes, their expression levels were not significantly different from those of microglia in the cortex. Under CPZ conditions, both cortical and CC microglia responded by increasing gene expression and significant DEGs (11% of genes detected in microglia in the cortex and CC regions) (Figure 5D). MOLs located in the CC and cortex had different signatures in the control condition, and both responded mildly to CPZ (3% of genes detected in MOLs in the cortex and CC regions) (Figure 5E). To highlight the regional cell type-specific responses to CPZ, we also analyzed astrocytes and OPCs. Cortical and CC astrocytes exhibited different expression signatures under the control condition, and cortical astrocytes exhibited a greater number of DEGs in response to CPZ (Figure S4B). OPCs located in the CC and cortex showed similar expression signatures, and they did not seem to respond to CPZ by changing their expression even though the number of OPCs markedly increased under CPZ conditions (Figure 5C and S4C).

### SpaTalk predicted heterogeneous cell–cell communication in the cortex and CC

We used SpaTalk to infer spatially resolved cell cell interactions from the deconvoluted Visium data ^43^ and focused on interactions among 4 cell types: microglia, astrocytes, the MOL, and the OPC. In the cortex, the interaction between astrocytes and microglia in the CPZ had the greatest number of identified receptor pathways, including death receptor, TNF, and interleukin family signaling pathways. The interaction between microglia and the MOL in the CPZ condition was associated with the second highest number of receptor pathways identified, including interleukin family signaling, ERK1/2 activation, and the Wnt signaling pathway. Most receptor pathways in the cortex were identified in the CPZ treatment group, while the MOL-to-OPC and OPC-to-MOL-enriched receptor pathways were identified in the control group (Table S1).

In the CC, fewer significant receptor pathways were identified than in the cortex, and astrocyte-to-microglia interactions continued to be associated with CPZ conditions. However, the interaction between the MOL and microglia had the greatest number of receptor pathways identified under all 3 experimental conditions, including PTK6, TNF, and CRMPs in Sema3A signaling. Microglia-MOL interaction receptor pathways were associated with RCV conditions (Table S2). Selected receptor pathways of interest from SpaTalk with a representative LR pair are plotted in Figure S5. Finally, we applied the same filters to CellChat and SpaTalk and identified 20 consensus LR pairs in the CC under CPZ conditions (Table 1). In 11 out of the 20 LR pairs, the LR pathway was enriched in growth factor and phagocytic pathways, and microglia were the major sender cells.

**Table 1.** Common LR pairs in CC under the CPZ condition predicted by both CellChat and SpaTalk.

| Sender | Receiver | Ligand | Receptor |
| --- | --- | --- | --- |
| astrocyte | microglia | Fgf1 | Fgfr2 |
| astrocyte | microglia | Gas6 | Axl |
| astrocyte | microglia | Pros1 | Axl |
| astrocyte | microglia | Vegfa | Flt1 |
| astrocyte | microglia | Vegfb | Flt1 |
| microglia | astrocyte | Fgf2 | Fgfr1 |
| microglia | astrocyte | Fgf2 | Fgfr2 |
| microglia | astrocyte | Fgf2 | Fgfr3 |
| microglia | astrocyte | Gas6 | Axl |
| microglia | astrocyte | Igf1 | Igf1r |
| microglia | astrocyte | Pdgfa | Pdgfrb |
| microglia | mol | Fgf2 | Fgfr2 |
| microglia | mol | Igf1 | Igf1r |
| microglia | opc | Fgf2 | Fgrf2 |
| microglia | opc | Igf1 | Igf1r |
| microglia | opc | Pdgfa | Pdgfra |
| mol | microglia | Gdf15 | Tgfbr2 |
| mol | microglia | Vegfa | Flt1 |

### The CPZ model exhibited moderate translatability to MS chronic active lesions

Finally, we aimed to assess the translatability of the CPZ model by performing an integrated analysis of our dataset with a published MS RNA-seq dataset that represents chronic active lesions ^7^. MS white matter lesions can be classified into various types based on the location of infiltrating immune cells, the activity of monocyte-derived macrophages and resident microglia, and the degree of demyelination ^20^. Among white matter lesions, disease disability correlates with chronic active lesions that have iron-laden microglia present at the lesion rim, which can be visualized by specific magnetic resonance imaging (MRI) sequences in the clinic ^7, 44^. We reanalyzed both human and mouse datasets using the same methods presented in this study and calculated DEGs between the disease and control groups to determine the level of DEG conservation between the two datasets. To take advantage of the different lesion regions defined by Absinta *et al*., we calculated DEGs of both “lesion core vs. control” and “chronic active lesion edge vs. control” for each glial cell type (Figure 6, left panel). We then calculated the overlap in DEGs between mice and humans and subsequently performed KEGG analysis on DEGs common to both species. We found pathways that were conserved between the CPZ mouse and human MS disease states, depending both on the cell type and the lesion architecture investigated (Figure 6, left panel).

**Figure 6.**
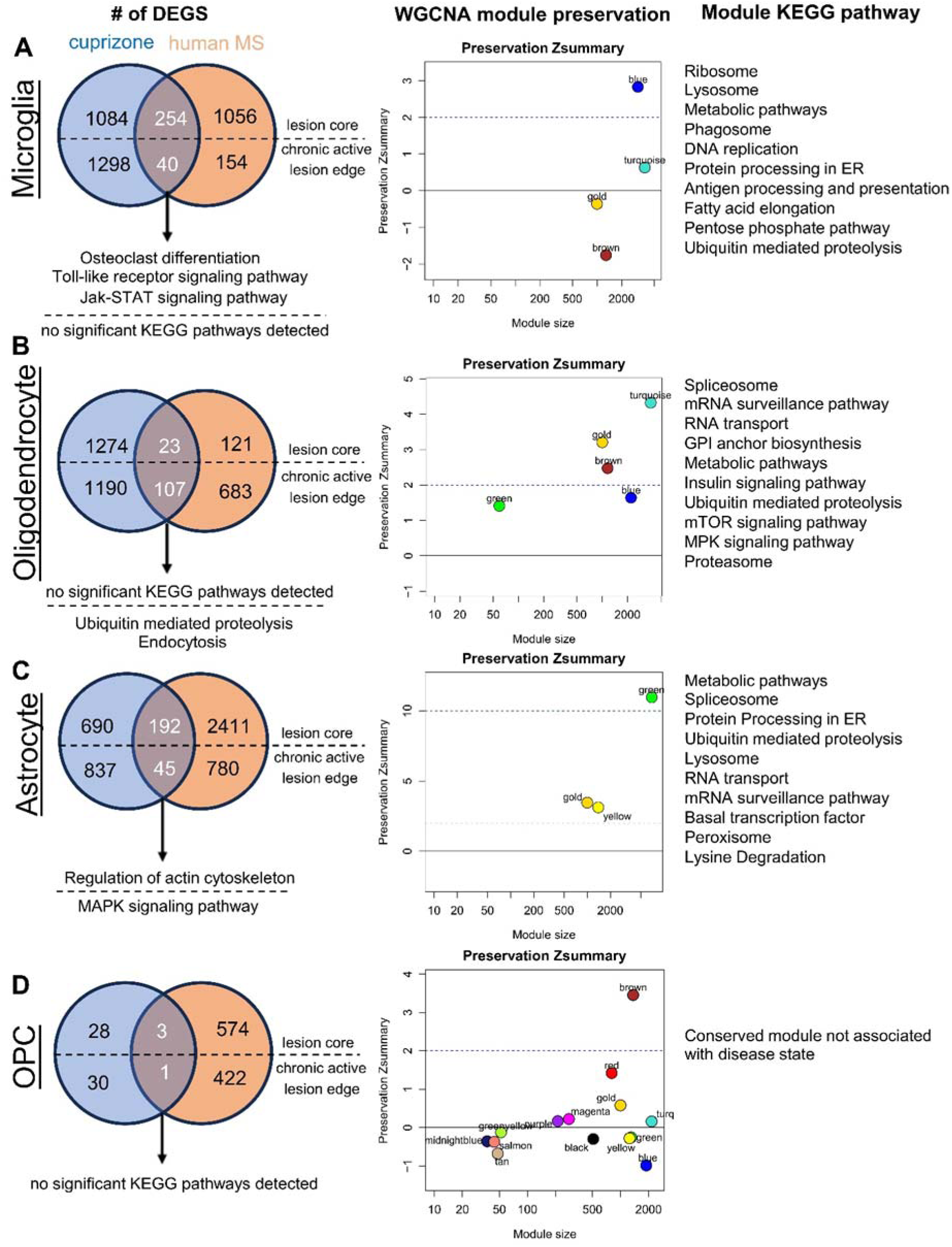
Translatability analysis revealed different levels of conservation of disease-associated modules driven by cell types between CPZ and human MS. (A) Comparison of DEG lists from CPZ (blue) and human MS (orange) in microglia (left). The number of genes shared between the two datasets is represented as white numbers at the intersection of the Venn diagram. DEGs from the CPZ were compared to those from two different lesion locations in the human data, either “control vs lesion core” (above dotted line) or “control vs. chronic active lesion edge” (below dotted line). KEGG pathway results using the shared genes as input are shown below each Venn diagram, with a dotted line again separating human comparisons of “lesion core” or “chronic active lesion edge.” hdWGCNA results showing the number of genes and preservation score of modules from microglia calculated from the CPZ data. A module preservation score >2 (dotted line) is considered moderately conserved between mice and humans, while a score > 10 is considered highly preserved. KEGGprofile results of the top preserved disease-associated module for microglia (right). (B) Comparison analysis of the MOL. (C) Comparison analysis of astrocytes. (D) Comparison analysis of OPCs. The moderately preserved brown module in the OPC cell type was not associated with disease and thus was not analyzed further.

We also used an alternative approach to investigate the translatability that is DEG agnostic by using a variation of weighted correlation network analysis designed for single-cell data (hdWGCNA) ^45, 46^. We used hdWGCNA to define disease-associated modules that were both correlated with the disease state and preserved between the two species ^47^. We found disease-associated modules in the CPZ model that showed moderate to strong preservation in humans depending on the cell type investigated (Figure 6, middle column). Astrocytes had the highest preservation scores of disease-associated modules, while MOL, OPC, and microglia showed moderate to low preservation between the CPZ and the Absinta *et al*. datasets. These disease-associated and preserved modules showed KEGG pathway enrichment that was shared among cell types, such as metabolic pathways (in astrocytes, microglia, and the MOL), protein processing in the endoplasmic reticulum (in astrocytes and microglia), and ubiquitin-mediated proteolysis (in astrocytes, microglia, and the MOL) (Figure 6, right column). In the future, this workflow can be applied to compare different MS lesion types to investigate the exact translatability of the CPZ model to MS pathology.

## Discussion

In this study, we profiled the CPZ model by “space” and “time”. We characterized the CPZ model by ST along with an array of bioanalytical tools to provide insights into biological processes during de and remyelination, as well as the translatability of the CPZ model to MS pathology. By ST, we found that demyelination and neuroinflammation in the CPZ model were more prominent and widespread across the dorsal to ventral axis than previously identified by IHC. Furthermore, the upregulation or downregulation of certain pathways, such as genes related to mitochondrial dysfunction and RhoA signaling, was cluster (brain region)-dependent based on the spatial resolution. We identified oligodendroglia and microglia as two major cell types associated with the CPZ model, and we identified the CPZ-associated subclusters of oligodendroglia and microglia located in the CC, the brain region of which shows stereotypical demyelination. In the RCV group, whereas the MOL normalized to the control state, the microglial phenotype continued to be associated with the CPZ, i.e., demyelination. We inferred cell cell interactions using both snRNA-seq and ST data and found that the phagocytosis, growth factor, and chemokine pathways were enriched under CPZ conditions.

ST is a fast-emerging field that enables high-throughput investigations of spatially resolved transcripts ^21, 22^. Prior to the development of ST technology, region-specific gene expression studies in the CNS relied heavily on manual dissections ^48^, which were sufficient to identify major biological function heterogeneity but lacked fine resolution. The key feature of MS diagnosis is tissue damage dissemination in space and time. To understand the pathology underlying MS disease activity and progression, MS lesions have been characterized extensively by both MR imaging and pathology examination ^2, 20^. At the tissue level, MS lesions exhibit heterogeneous architectures and cell type involvement; therefore, MS lesions are often manually dissected for gene profiling to better understand disease mechanisms ^7, 9, 49^. Based on our study of the CPZ model, the ST appears to be a superior tool for investigating the detailed cellular and molecular substrates underlying MS lesion evolution to manual dissection.

Microglia and oligodendroglia are 2 major cell types significantly associated with the CPZ model, and their subclusters associated with the CPZ phenotype are consistent with what has been reported previously ^19^. Our findings are consistent with the IHC results showing that demyelination (oligodendrocytes) and gliosis (microglia) are the two prominent features of the CPZ model, particularly in CC lesions ^25^. However, considering the proposed functions of astrocytes in the experimental autoimmune encephalomyelitis (EAE) model and MS ^50, 51, 52^, it is unexpected that astrocytes were not significantly associated with any experimental conditions, and no disease-associated subclusters were identified in our dataset. It is possible that (1) the recovery time point (2 weeks after CPZ withdrawal) is too early to capture a full astrocytic response in our CPZ model or (2) astrocyte involvement in demyelination pathology is animal model dependent, as the EAE model is more inflammatory in nature. Nonetheless, CellChat and SpaTalk predicted numerous potential LR pairs between astrocytes and microglia, highlighting the interactions between the 2 cell types in the CPZ and the neuroinflammatory pathways associated with these interactions. Interestingly, astrocytes had the highest preservation scores of disease-associated modules with MS among all the cell types analyzed by hdWGCNA for model translatability. Although we did not identify astrocytes as “pathogenic” cells in the CPZ model, this finding suggested that astrocyte transcriptomic changes were still in line with the changes in MS.

Our analysis indicated that in the RCV group, whereas the MOL normalized to the control state, the microglial phenotype continued to be associated with the demyelination phenotype. This finding is consistent with studies using IHC as their major readout to show robust spontaneous remyelination in the model. A complete functional remyelination phenotype is thought to consist of the introduction of OPCs to the lesion, the differentiation of OPCs into the MOL and finally myelin compaction. Although the expression signature in RCV oligodendroglia was normalized to that in control cells, our study did not investigate the functional normalization of MOLs. It is possible that part of the expression signature is driven by a cell intrinsic differentiation program and that full remyelination/recovery requires further interactions with other cell types, such as microglia ^53^.

The growth factor and phagocytic pathways were predicted by both CellChat and SpaTalk as potentially significant LR interactions in the CC under CPZ conditions. The role of growth factors in MS has long been recognized ^54^, and approximately three-quarters of the LR pairs are related to growth factor signaling. There are 18 members of the fibroblast growth factor (FGF) family. Through FGFR signaling, FGFs are involved in various biological and pathological processes, and FGF1 and FGF2 are the main mitogens in the CNS ^55^. Evidence has shown that FGF1 and FGF2 are expressed in MS lesions with a remyelination phenotype ^56, 57^, and it has also been shown that FGF signaling is critical for regulating oligodendrocyte and myelin regeneration ^58^. Thus, recombinant FGFs have been proposed to be a potential therapeutic approach for MS. On the other hand, oligodendrocyte-specific deletion of FGFR1 and FGFR2 results in a protective effect in the EAE model; therefore, FGFR inhibitors have also been proposed for MS ^59^. Disentangling the complex biology of FGF signaling will be critical for MS therapy development. The remaining LR pairs are related to phagocytic pathway signaling. Myelin debris has been shown to inhibit remyelination ^60^, and it was further suggested that a dysfunctional myelin sheath under inflammatory attack could pose a greater risk of axonal degeneration ^61^. Therefore, a precise and efficient phagocytic program to clean myelin debris is critical for successful remyelination. In nondemented individuals from the Swedish BioFINDER-2 cohort, increased levels of microglial markers in the CSF, such as soluble TREM2, AXL, MERTK, GAS6, LPL, CST7, SPP1 and CSF1, are associated with slower cognitive decline ^62^. In our dataset, Gas6-Axl and Pros1-Axl were in the consensus LR pair list from CellChat and SpaTalk, while Gas6-Mertk and ApoE-Trem2 were predicted from the CellChat analysis. Gas6 and Pros1 are ligands for the TAM (Tyro3, Axl, and Mertk) receptor tyrosine kinase family, and LR signaling regulates inflammation, cell growth, and clearance of debris ^63^. The role of phagocytosis-related proteins in MS and whether they can be potential therapeutic targets are less clear than those in AD research ^64^; however, studies in the CPZ model have suggested that both Mertk and Trem2 play roles in myelin debris clearance and subsequently promote remyelination ^35, 65^. A detailed profiling of phagocytic proteins in MS CSF to see how they correlate with MS disease severity and progression could be a worthwhile effort.

Overall, the LR pairs predicted in the CPZ condition in our study showed similarities to cell cell interactions identified from human MS cortical ST data ^66^. For example, growth factor pathway interactions, such as those involving PSAP-GPR37L1, CSF1-CSF1R, and PDGFC-PDGFRA, which represent approximately one-third of the pathways identified in the MS cortical ST data, were also present in our CPZ dataset. One-fourth of the pathways in the MS cortical ST data were grouped as anti-inflammatory, such as the PROS1-AXL interaction, which was also predicted from the CPZ dataset. On the other hand, the complement pathway, which has been shown to be critical in MS pathology ^7, 66^, was not one of the top pathways identified in our study. This suggests that it is important to first understand what aspects of the disease can be modeled by animal studies. We further demonstrated that the translatability could be cell type dependent. That is, astrocytes had the highest preservation scores of disease-associated modules, while MOL, OPC, and microglia showed moderate to low preservation scores between the CPZ model and MS chronic active lesions. To fully utilize the CPZ model, we suggest that transcriptomic analysis should also be performed when possible, in addition to IHC.

The advantage of our study is that we employed a hypothesis-free whole-genome approach, but we also encountered some study limitations. First, while cell cell interaction investigations in our study provided novel findings, the LR pairs did not immediately predict therapeutic directionalities. Due to the scope of the study, functional validation will be performed in a separate investigation. Second, it is important to acknowledge that 10X Visium technology lacks single-cell resolution, which is a key limitation. To address this issue, we used cell2location, which integrates snRNA-seq data with spatial data to deconvolute cell types, providing insights into the abundances of different cell types within the 10X Visium spots. Although this technique does not achieve true single-cell resolution, it represents a crucial step toward a more detailed characterization of the cellular landscape. Finally, while we believe that the CPZ model captures some of the salient demyelination features of MS, we cannot exclude direct effects of CPZ on cells other than the oligodendrocyte population or effects from CPZ on biological processes unrelated to demyelination, which could introduce noise into the analyses ^67^.

## Materials and Methods

### Cuprizone model

All procedures involving animals were reviewed and approved by the Biogen Institutional Animal Care and Use Committee and conducted under protocol 801 in accordance with relevant guidelines and regulations. All methods are reported in accordance with ARRIVE guidelines (https://arriveguidelines.org). Male wild-type C57BL/6J mice aged between 8 and 10 weeks were purchased from Charles River Laboratories (stock number 027C57BL/6). The research animals were housed under specific-pathogen-free (SPF) conditions in an AAALAC accredited facility according to Biogen’s institutional animal care and use committee (IACUC) protocol with a 12 hr–12 hr light–dark cycle and environmental conditions controlled at 72°F/22°C and 40–60% humidity. To establish the CPZ model, customized 0.3% (w/w) cuprizone chows were made by mixing cuprizone powder (Sigma Aldrich 14690) with control base chows (AIN-76A) at Research Diets Inc. All food chows (CPZ and control) were stored in vacuum bags at 4°C, and fresh food chows were replaced weekly during the study. In the CPZ group, the animals were fed cuprizone chow for 4 weeks. For the recovery group, animals were fed cuprizone chows for 4 weeks, followed by 2 weeks of control base chows for recovery. For the control group, animals were provided with control base chow for 6 weeks. The body weight of each mouse was closely monitored weekly throughout the study, and mice with ≥ 30% weight loss were excluded.

### Tissue collection for sequencing studies

For snRNA-seq, spatial transcriptional sequencing, and bulk RNA-seq studies, the CPZ-treated mice were sacrificed, and brain tissues were collected at either week 4 (n=3, CPZ group) or week 6 (n=3, recovery group). The control mouse brain tissues were collected at week 6 (n=4, control group). At each time point, the mice were euthanized by CO_2_ inhalation, followed by transcardiac PBS perfusion and brain tissue dissection. Each brain was cut sagittally to separate the left and right hemispheres; one brain hemisphere was trimmed into three coronal segments and embedded in OCT for snRNA-seq or spatial transcriptomic study, and the other hemisphere was snap frozen for bulk RNA-seq analysis (Figure 1). To enrich for cells from the corpus collosum and cortex, only the top half of the coronal section, which included the dorsal cortex, CC, and hippocampus, was collected for snRNA-seq.

### Immunohistochemistry (IHC)

A separate cohort of animals was used for the IHC study. Brain tissues were dissected and fixed for 24 hours in 10% NBF at 4°C and then transferred to ice-cold PBS for coronal sectioning on a vibratome. Coronal brain slices of 40 µm thickness from bregma –1 mm to –2.5 mm were collected and stored in 30% sucrose/PBS buffer at -20°C. For staining, 3 slices spanning 400 µm apart were selected from each animal. For staining, brain slices were thawed in PBS at room temperature and then blocked with blocking solution (0.5% Triton, 5% goat serum, and 0.1% BSA in PBS) for 1 hour at room temperature. Primary antibodies were diluted in blocking solution and incubated with the slices overnight at 4°C. The corresponding secondary antibodies were applied for 1-2 hours at room temperature. After incubation with secondary antibodies, the slices were washed and mounted onto glass coverslips with Prolonged Gold Mounting Solution (Thermo Fisher, P36931). The following primary antibodies were used: mouse anti-MBP (BioLegend, 88408, 1:500), rat anti-CD68 (Bio-Rad, MCA1957, 1:300), and chicken anti-GFAP (Thermo Fisher, PA1-10004, 1:500). The following secondary antibodies were used: goat anti-chicken IgY 647 (Thermo Fisher, A32933, 1:500) and donkey anti-rat IgG488 (Thermo Fisher, A21208, 1:500).

Images were obtained using a Zeiss LSM 710 confocal microscope with a 20x objective and Z-stack of 10 planes at 1 µm intervals. Individual fields of each slice were tiled into a large image.

### Tissue sectioning and preparation for spatial transcriptomics

Frozen sections for ST profiling were collected using a Leica CM1520 cryostat with a Leica Surgipath high profile blade at -23°C. Ten-micron-thick sections were collected from mouse brain hemispheres embedded in OCT (TissueTek) on Visium Spatial Gene Expression slides (10X Genomics Slide Kit, Ref No. 1000185). Tissue sections were carefully collected to allow accurate positioning within individual capture areas of Visium slides following a randomized collection scheme across multiple Visium slides to help control for (1) slide-to-slide batch effects by ensuring that a section from each treatment group was present on each slide and (2) the potential impact section location on the slide by ensuring that sections from a given treatment group were not in the same position or order on every slide. The frozen slides were processed for spatial transcriptomic sequencing according to the manufacturer’s instructions. Briefly, the sections were heated at 37°C for 1 minute and then fixed in methanol at –20°C for 30 minutes.

Following fixation, the samples were stained with hematoxylin and eosin. Brightfield images were collected on an Olympus VS120 Virtual Slide Microscope equipped with a VC50 camera using a UPLSAPO 20X objective. After imaging, tissue permeabilization was carried out using the supplied permeabilization buffer at 37°C for 6 minutes. Permeabilization time was selected based on a prior optimization experiment in mouse brain tissues using Visium Spatial Tissue Optimization slides (10X Genomics, Ref. No. 1000193). Following permeabilization, reverse transcription and second strand synthesis reactions were carried out on slides before elution of the resulting cDNA into separate sample PCR tubes according to Visium guidelines. qPCR was performed on 1 µL of eluted cDNA as described in the Visium Spatial Gene Expression Reagent Kits User Guide (Revision-B) and was run on an Applied Biosystems QuantStudio 12K Flex qPCR instrument (Applied Biosystems™ 4471087). Individual sample cDNA PCR amplification cycles were selected and run based on sample cDNA Ct. Total yields of amplified cDNA ranged from 300-550 ng. Sequencing library generation was performed using the 10X Genomics Library Construction Kit (PN-1000190) following the manufacturer’s instructions with 13 cycles of PCR based on the cDNA input used. Library quality was measured using a PerkinElmer LabChip GXII with a Hi Sens Lab Chip (Perkin Elmer 760517) and DNA High Sensitivity Reagents (Perkin Elmer CLS760672), which showed an average spatial gene expression library size of 473 bp. Libraries were normalized and pooled before loading at 300 pM on an Illumina NovaSeq 6000 S2 flow cell with paired-end sequencing parameters of Read1-28 bp x i7-10 bp x i5-10 bp x Read2-120 bp.

### Bulk RNA sequencing and data processing

Bulk RNA-seq libraries were prepared using the Kapa mRNA HyperPrep Kit (Kapa Biosystems KK8581) according to the manufacturer’s instructions. Briefly, 100 ng of RNA was fragmented for 6 min at 94°C, followed by RT, second strand synthesis, A-tailing, adaptor ligation, and amplification with 14 cycles of PCR. Libraries were quantified on a LabChip GXII as described above. Libraries were normalized, pooled, and sequenced on an Illumina HiSeq2500 platform, with an average depth of 9 million 50 bp paired-end reads. Run quality metrics were assessed using Illumina’s BaseSpace run summary tool. Bulk RNA analysis was performed using the RNASequest pipeline ^68^ following the tutorial provided in the manuscript.

### Bulk RNA-Seq integration with scRNA-Seq

We applied Scissor ^32^ (https://github.com/sunduanchen/Scissor) to integrate the bulk and snRNA-Seq datasets. It utilizes the phenotypes (CPZ, RCV, and control) collected from bulk assays to identify the most highly phenotype-associated cell subpopulations from single-cell data.

### Nuclei Isolation

Nuclei isolation was performed as described previously ^69^. Tissue was dounced in 0.5 mL Hyman Lysis Buffer (0.32 M Sucrose, 5 mM CaCl_2_, 3.0 mM MgAc_2_, 0.1 mM EDTA, 10 mM Tris-HCl, pH 8.0, 1 mM DTT, 0.1% Triton X-100, 1/1000th volume Promega RNasin Plus RNase Inhibitor) 10X with loose pestle, followed by 10X with tight pestle. Following douncing, 1 mL of Hyman Low Sucrose Buffer (0.32 M Sucrose, 5 mM CaCl_2_, 3.0 mM MgAc_2_, 0.1 mM EDTA, 10 mM Tris-HCl, pH 8.0, 1 mM DTT, 1/1000th volume Promega RNasin Plus RNase Inhibitor) was added to the homogenate, and passed through a 70 μm filter, followed by a 1 mL Hyman Low Sucrose Buffer rinse of the dounce and filter. Homogenates were spun at 600xg for 5-10 min at 4°C. After supernatant aspiration, pellets were gently resuspended in 5 mL Complete Buffer HB (0.25 M sucrose, 25 mM KCl, 5 mM MgCl_2_, 20 mM Tricine-KOH, pH 7.8, 1 mM DTT, 0.15 mM spermine, 0.5 mM spermidine, 1/500th volume Promega RNasin Plus RNase Inhibitor), followed by addition of 5 mL Working Solution (50% Iodixanol: 5 volumes OptiPrep Density Gradient Medium, 1 volume Diluent Buffer (150 mM KCl, 30 mM MgCl_2_, 120 mM Tricine-KOH, pH 7.8), 1/1000th volume Promega RNasin Plus RNase Inhibitor). In a 12 mL Corex tube, 1 mL 40% iodixanol (Working Solution diluted in Complete Buffer HB, 0.042% BSA), and 30% iodixanol (Working Solution diluted in Complete Buffer HB, 0.021% BSA) were layered, followed by 10 mL of resuspended sample. The samples were spun in an ultracentrifuge at 10,000xg for 18 min at 4°C with the brake off. After spinning, the fatty myelin top layers were vacuum aspirated, followed by a 25% iodixanol layer and approximately 1/3 of the 30% iodixanol layer. Approximately 0.5 ml of nuclear sample band at the 30%-40% iodixanol interface was collected, transferred to an Eppendorf tube and mixed by pipetting. Nuclei were stained with DAPI and counted with a Countess II FL. Nuclei suspensions were diluted to 1000 nuclei/μL in RB (PBS + 2% BSA + 1/1000th volume Promega RNasin Plus RNase Inhibitor).

### snRNA sequencing

Single-nuclei RNA-seq was performed using the 10X Genomics Chromium Next GEM Single Cell 3’ Kit v3.1 (PN1000268), Chromium Next GEM Chip G (PN1000127), and Dual Index Kit TT Set A (PN1000215). Nuclei suspensions were loaded on the 10X Chromium instrument for recovery of 5000 nuclei per sample. cDNA synthesis, PCR, and library preparation were performed according to the manufacturer’s instructions, with 12 cycles of PCR for cDNA amplification, and 15 cycles for library amplification. cDNA and libraries were quantified on a LabChip GXII as described above. Libraries were normalized, pooled, and loaded at 300pM on an Illumina NovaSeq 6000 S4 flowcell with paired end sequencing parameters of Read1-28bp x i7-10bp x i5-10bp x Read2-90bp targeting an average depth of 85,000 reads per cell.

### snRNA-seq data preprocessing and quality control

FASTQ files were generated from Illumina bcl files using CellRanger mkfastq. CellRanger count was used to identify UMI and cell barcodes and to align reads to the mouse genome assembly mm10, with intronic counts included. To remove ambient RNA from the gene expression matrices, the ‘remove-background’ function of CellBender was used ^70, 71^. In addition, CellBender identifies and removes empty droplets. The CellRanger output file ‘raw_feature_bc_matrix.h5’ was used as the input file for CellBender. The ‘Estimated Number of Cells’ from CellRanger was used as the ‘expected-cells’ parameter, while a value in the plateau of the barcode rank plot was used as the ‘total-droplets-included’ parameter. Potential doublets were identified and removed using scDblFinder ^72^. Filtered barcode matrices were concatenated using Scanpy ^73^. QC metrics, including the number of detected genes, total counts, and percentage of counts from mitochondrial genes, were calculated for each barcode. Cell barcodes with fewer than 1000 detected genes, greater than 100000 counts, or greater than 10% mitochondrial UMIs were filtered out from the dataset.

As an additional QC metric, intronic mapping rates were derived on a per-barcode basis using the BAM alignment files output by CellRanger count. Compared with cellular transcriptomes, nuclear transcriptomes have been demonstrated to show a greater proportion of reads mapping to intronic regions ^74^, enabling the detection of nonnuclear cell barcodes representing contaminating cytoplasmic transcripts.

### snRNAseq cell type annotation and subclustering

Cell type labels were assigned using label transfer from E-MTAB-11115 ^29^, with region-specific astrocyte subtypes merged into a single astrocyte label. Oligodendrocyte-lineage cell types were further clarified by label transfer from GSE75330 ^30^ after merging major cell types from this reference. Filtered UMI counts for E-MTAB-11115 were downloaded from ArrayExpress along with cell type annotations. The UMI was then processed by Seurat ^75^, LIGER ^76^ and Harmony ^77^ integration on different sample libraries. The kBET ^78^ evaluation revealed that LIGER achieved the best performance. Thus, the Seurat SCT and LIGER layout (PCA) was used to generate the Seurat azimuth reference. In parallel, the cells were sequentially subclustered via Harmony and Louvain clustering.

To investigate the heterogeneity within the microglial and oligodendrocyte populations, we subclustered these cell types using the Leiden clustering algorithm implemented in Scanpy. By applying a resolution parameter of 0.5, we identified 12 subclusters each for microglia and oligodendrocytes. Differential gene expression analysis was performed to identify marker genes that exhibited significant differences in expression between subclusters with the Wilcoxon rank-sum test. To visualize marker gene expression, we generated dot plots that depict the percentage of cells expressing the gene through dot size, and the color intensity represents the relative expression level.

### snRNA-seq DE Analyses

snRNA-seq data were analyzed per cell type using the NEBULA-HL single cell DE method ^79^. For each cell type, 3 contrasts were considered: CPZ vs Control, CPZ vs Recovery, and Recovery vs Control. Cell types with at least 10 cells in each of the biological replicates within the contrast were analyzed. The genes tested in each contrast group were expressed in at least 10% of the cells in both contrast groups. Mouse ID was used as a random effect in the model. Library size was used as an offset covariate.

### Pseudotime trajectory analysis and velocity visualization

We used scFates ^80^ for trajectory analysis. scFates combines a tree inference approach similar to SimplePPT ^81^ and a principal graph learning algorithm named ElPiGraph ^82^ to calculate trajectories.

### 10x Visium spatial sequencing data analysis

**The** 10x Visium spatial sequencing data were processed using Space Ranger v1.1.0 (10x Genomics). The sequencing results (BCL files) were converted to fastq files with the Space Ranger mkfastq pipeline. The fastq files were then mapped to the mouse reference genome (GRCm38.p3) using the Space Ranger count pipeline, which produces a spot-barcoded expression matrix.

Next, we used SpaGCN ^23^, a graph convolutional network-based method, to identify spatial clusters based on both gene expression profiles and spatial locations. All the slides were aligned and normalized using the PASTE algorithm ^83^. The MS gene signature included multiple sclerosis signaling, myelination signaling and neuroinflammation signaling pathways: B2m, C1qa, C1qb, C1qc, C3, C3ar1, C4b, Ccl2, Cd86, Creb3l1, Cxcl10, Egr2, Eif4ebp1, Fos, Hck, H2-Q4, H2-Dmb1, H2-Oa, H2-Aa, H2-Ab1, H2-Eb1, Hmox1, Icam1, Idi1, Igf1, Il33, Irf7, Itgax, Itgb2, Itgb4, Itgb5, Mag, Mapk4, Mbp, Mobp, Mog, Mthfd2, Ncf2, Opalin, Parp14, Parp9, Plp1, Pycard, Rac2, Rnf213, Serping1, Slc6a12, Tgfbr1, Tlr2, Tlr3, Tlr7, Trem2, Tyrobp.

Then, we used Cell2location ^29^ to spatially map cell types in the Visium spatial transcriptomics data. This package integrates single-cell RNA-seq (scRNA-seq) or single-nucleus RNA-seq (snRNA-seq) with spatial transcriptomics data to deconvolute cell types in each spot of the Visium slide. A reference mouse brain cell type signature representing gene expression profiles for each annotated cell type was estimated from snRNA-seq data using a negative binomial regression model. The parameters used for the model were batch size=2500, train_size=1, learning rate=0.002, and max_epochs=250. Then, the reference cell type signature was integrated with the processed 10X Visium data to deconvolute the absolute spatial abundance of each cell type on the spatial transcriptomics slide. The deconvolution model requires the user to input two hyperparameters: 1) the expected cell abundance (N_cells_per_location), which was set to 5, and 2) the regularization strength of the detection efficiency effect (detection_alpha), which was set to 200 (default). The model was trained for 30,000 iterations. The default settings were used for the remaining parameters.

### ST pseudobulk PCA

To perform pseudobulk PCA of the spatial transcriptomics data, spot counts for each gene were aggregated across all spots within each slice to form a gene-by-slice count matrix. After performing variance stabilizing transformation (vst), the top 500 most variable genes from the normalized matrix were used to generate PCs. The plot shown in Figure 1B was generated from data processed with DESeq2 ^84^ after correcting for batch effects using ComBat-seq ^85^.

### ST DE and pathway analysis

ST spot-level data were analyzed per cell type using the NEBULA-HL single-cell DE method ^79^. In this analysis, each spot was handled independently (spot-level information regarding spatial adjacency was not considered). Regional subsets of mouse brain regions, including the CC, cortex, and hippocampus, were manually annotated. For each region, 3 contrasts were considered for DE analysis, namely, CPZ vs Control, CPZ vs RCV, and RCV vs Control. Spots with fewer than 100 counts or greater than 45% mitochondrial gene counts were filtered out from each slice before analysis. Mouse ID was used as a random effect in the model. Library size was used as an offset covariate. DEGs from each cluster were analyzed by QIAGEN Ingenuity Pathway Analysis (IPA) for pathway discovery using Core Analysis. To compare pathways across clusters, all 4 clusters were subjected to Comparison Analysis.

### Cell–cell interaction analysis

We applied CellChat ^34^ to understand the cell-cell interactions from the snRNAseq dataset. For ligand-receptor (LR) analysis, the scRNA-seq count matrix and corresponding metadata were loaded into a Seurat object for use in CellChat. The count matrix was normalized, and cell type identities were assigned to cells. The mouse specific CellChat LR interaction database ‘Secreted Signaling’ was selected. CellChat with default settings was used to perform cell-cell interaction (CCI) analysis and infer the CCI network by computing cellular and signaling pathway communication probabilities (strength). The method truncatedMean was used to calculate the average gene expression per cell group. Each comparison (CPZ vs. Control, CPZ vs. Recovery, Recovery vs. Control) was analyzed separately and then merged into a single CellChat object. Differential LR combinations within the CCI network were identified in CellChat by comparison of the communication probabilities, and then those results were extracted for the cell type pairs of interest.

We applied SpaTalk ^43^ to infer cell cell interactions by enriching receptor pathways from the ST dataset. SpaTalk was run on three spatial sections per condition, with one section per biological replicate. Scaled cell type deconvolution weights precalculated by cell2location were used to deconvolve spots into cell types that were derived from the scRNA-seq reference data. Deconvoluted spots from two annotated regions, the cortex and CC, were analyzed in each section. Cell type assignments per spot were made using the default recommended size for 10X Visium 55-micron spots. Default SpaTalk spot-level analysis parameters were used to calculate cell cell LR interactions and to perform enriched LR receptor pathway analysis. Overlapping significant cell cell LR interaction pairs from both CellChat analysis based on snRNA-seq data and SpaTalk analysis based on ST data were used for downstream comparisons between CPZ and control conditions. The following set of 8 custom LR interaction pairs relevant to MS were added to the overlapping set: Il34-Csf1r, Apoe-Trem2, Pecam1-Pdgfra, Th-Drd1, Th-Drd2, Ddc-Drd1, Ddc-Drd2, and Spp1-Cd44.

### Translatability and hdWGCNA

For comparative DEG analyses between CPZ and the human dataset, we first reprocessed the data from Absinta *et al*. ^7^ (see snRNA-seq data preprocessing and quality control) and called DEGs between control and MS tissues using Nebula (see snRNA-seq DE Analyses). For the human dataset, we performed DEG analysis on both the “control vs lesion core” and control vs. “chronic active lesion edge” datasets to take advantage of the different lesion areas provided by Absinta *et al*. We identified genes that were upregulated in the disease state, with an FDR < 0.05 for both datasets. We then converted mouse gene IDs into their human orthologs and counted the number of distinct and overlapping DEGs between the two datasets on a cell type basis. For a given cell type, DEGs that were common between CPZ and humans were used as input to the R package KEGGprofile for pathway analysis (https://github.com/slzhao/KEGGprofile/blob/master/DESCRIPTION).

Single-cell WGCNA was performed according to hdWGCNA ^45, 46^ with modifications to regress out pseudobulk covariates, such as age and sex. The working principle of hdWGCNA is to first create pseudobulk samples of single-nucleus data based on kNN and subsequently identify groups of coexpressed genes (“modules”) for each cell type. Briefly, raw pseudobulk expression was calculated from specific cell populations by aggregating UMI counts from 50 cells using a k-nearest neighbor approach. We then normalized the raw count data by variance stabilizing transformation (VST) implemented by DESeq2. We regressed out confounding features such as total UMI count, mitochondrial percentage, and intronic rate. We then applied WGCNA to identify coexpressed genes (modules) using the normalized count data. Briefly, WGCNA starts by constructing a topological overlap (TO) matrix ^86^. To this end, we used weighted correlation with individual sample weights determined as described above and the “signed hybrid” network in which negatively correlated genes are considered unconnected. We used module eigengenes (MEs), the first principal component of the module’s gene expression matrix, to represent the overall expression patterns of the corresponding coexpression modules. The disease correlation score was defined as the Pearson correlation coefficient between the module eigengenes and disease status. Finally, we performed module preservation analysis using the modulePreservation function in WGCNA. The level of translatability between species is defined by the module preservation score ^47^. The output consists of “modules” of genes with similar trajectories, including scores to assess both correlation with disease (module correlation) and preservation between mice and humans (Zsummary). We considered modules with a disease correlation score >0.7 and a Zsummary >2 to be associated with disease and preserved between species. For pathway analysis, we used the module as an input for the KEGGProfiler.

## Data Availability

The raw sequencing data associated with this study are available through the Gene Expression Omnibus (GEO) database. The superseries GSE255371 (https://www.ncbi.nlm.nih.gov/geo/query/acc.cgi?acc=GSE255371) can be used to access the raw data. These superseries include bulk RNA-Seq (GSE255368), snRNA-Seq (GSE255369) and spatial transcriptomics (GSE255370) datasets. All analysis codes are available in the GitHub repository: https://github.com/interactivereport/Mouse_Cuprizone_Spatial.

## Author Contributions

HHT, AJG, TMC, BZ and KL designed and implemented the study and analysis plan. JW, AJG, SJC, PC, RC, WZ, YJK-W, CE, LJ, HM, and TMC were involved in sample preparation, the Visium experiment, image scanning and NGS data generation. AJG, SJC, and TLR performed the pathology review. SP, WH, JZ, ARG, SC, MS, JC, ZO, MR, ML, WW, EZ, EM, JG, and ML performed the data analysis. HHT, SP, and BZ wrote the manuscript with input from all the authors. All the authors have read and approved the manuscript.

## Acknowledgment

The authors thank Drs. Bing Zhu and Robert Miller for critically reading the manuscript and for discussions with team members.

## References

1. Walton C, et al. Rising prevalence of multiple sclerosis worldwide: Insights from the Atlas of MS, third edition. *Multiple sclerosis (Houndmills, Basingstoke*, England) 26, 1816–1821 (2020).

2. Reich DS, Lucchinetti CF, Calabresi PA. Multiple Sclerosis. N Engl J Med 378, 169–180 (2018).

3. Lassmann H. Multiple Sclerosis Pathology. Cold Spring Harb Perspect Med 8, (2018).

4. Klotz L, Antel J, Kuhlmann T. Inflammation in multiple sclerosis: consequences for remyelination and disease progression. Nature reviews Neurology 19, 305–320 (2023).

5. International Multiple Sclerosis Genetics C, Multiple MSC. Locus for severity implicates CNS resilience in progression of multiple sclerosis. Nature, (2023).

6. Consortium IMSG. Multiple sclerosis genomic map implicates peripheral immune cells and microglia in susceptibility. Science 365, eaav7188 (2019).

7. Absinta M, et al. A lymphocyte-microglia-astrocyte axis in chronic active multiple sclerosis. Nature, (2021).

8. Schirmer L, et al. Neuronal vulnerability and multilineage diversity in multiple sclerosis. Nature, (2019).

9. Jäkel S, et al. Altered human oligodendrocyte heterogeneity in multiple sclerosis. Nature, (2019).

10. Vega-Riquer JM, Mendez-Victoriano G, Morales-Luckie RA, Gonzalez-Perez O. Five Decades of Cuprizone, an Updated Model to Replicate Demyelinating Diseases. Curr Neuropharmacol 17, 129–141 (2019).

11. Narine M, Colognato H. Current Insights Into Oligodendrocyte Metabolism and Its Power to Sculpt the Myelin Landscape. Front Cell Neurosci 16, 892968 (2022).

12. Blakemore WF. Observations on oligodendrocyte degeneration, the resolution of status spongiosus and remyelination in cuprizone intoxication in mice. J Neurocytol 1, 413–426 (1972).

13. Barkhof F, et al. Remyelinated Lesions in Multiple Sclerosis: Magnetic Resonance Image Appearance. Archives of Neurology 60, 1073–1081 (2003).

14. Ransohoff RM. Animal models of multiple sclerosis: the good, the bad and the bottom line. Nat Neurosci 15, 1074–1077 (2012).

15. Gudi V, Gingele S, Skripuletz T, Stangel M. Glial response during cuprizone-induced de- and remyelination in the CNS: lessons learned. Front Cell Neurosci 8, 73 (2014).

16. Gudi V, et al. Synaptophysin Is a Reliable Marker for Axonal Damage. J Neuropathol Exp Neurol 76, 109–125 (2017).

17. Stidworthy MF, Genoud S, Suter U, Mantei N, Franklin RJ. Quantifying the early stages of remyelination following cuprizone-induced demyelination. Brain Pathol 13, 329–339 (2003).

18. Masuda T, et al. Spatial and temporal heterogeneity of mouse and human microglia at single-cell resolution. Nature, (2019).

19. Hou J, et al. Transcriptomic atlas and interaction networks of brain cells in mouse CNS demyelination and remyelination. Cell reports 42, 112293 (2023).

20. Kuhlmann T, Ludwin S, Prat A, Antel J, Bruck W, Lassmann H. An updated histological classification system for multiple sclerosis lesions. Acta Neuropathol 133, 13–24 (2017).

21. Rao A, Barkley D, Franca GS, Yanai I. Exploring tissue architecture using spatial transcriptomics. Nature 596, 211–220 (2021).

22. Liu B, Li Y, Zhang L. Analysis and Visualization of Spatial Transcriptomic Data. Frontiers in genetics 12, 785290 (2021).

23. Hu J, et al. SpaGCN: Integrating gene expression, spatial location and histology to identify spatial domains and spatially variable genes by graph convolutional network. Nat Methods 18, 1342–1351 (2021).

24. Xie M. Rostrocaudal Analysis of Corpus Callosum Demyelination and Axon Damage Across Disease Stages Refines Diffusion Tensor Imaging Correlations With Pathological Features. J Neuropathol Exp Neurol, (2010).

25. Praet J, Guglielmetti C, Berneman Z, Van der Linden A, Ponsaerts P. Cellular and molecular neuropathology of the cuprizone mouse model: clinical relevance for multiple sclerosis. Neuroscience and biobehavioral reviews 47, 485–505 (2014).

26. Orthmann-Murphy J, et al. Remyelination alters the pattern of myelin in the cerebral cortex. eLife 9, (2020).

27. Itoh N, et al. Cell-specific and region-specific transcriptomics in the multiple sclerosis model: Focus on astrocytes. Proc Natl Acad Sci U S A, (2017).

28. Harrington EP, Zhao C, Fancy SP, Kaing S, Franklin RJ, Rowitch DH. Oligodendrocyte PTEN is required for myelin and axonal integrity, not remyelination. Ann Neurol 68, 703–716 (2010).

29. Kleshchevnikov V, et al. Cell2location maps fine-grained cell types in spatial transcriptomics. Nat Biotechnol, (2022).

30. Marques S, et al. Oligodendrocyte heterogeneity in the mouse juvenile and adult central nervous system. Science 352, 1326–1329 (2016).

31. Scheyltjens I, et al. Single-cell RNA and protein profiling of immune cells from the mouse brain and its border tissues. Nat Protoc 17, 2354–2388 (2022).

32. Sun D, et al. Identifying phenotype-associated subpopulations by integrating bulk and single-cell sequencing data. Nat Biotechnol 40, 527–538 (2022).

33. Keren-Shaul H, et al. A Unique Microglia Type Associated with Restricting Development of Alzheimer’s Disease. Cell 169, 1276–1290 e1217 (2017).

34. Jin S, et al. Inference and analysis of cell-cell communication using CellChat. Nature communications 12, 1088 (2021).

35. Shen K, et al. Multiple sclerosis risk gene Mertk is required for microglial activation and subsequent remyelination. Cell reports 34, 108835 (2021).

36. Thurston G, Daly C. The complex role of angiopoietin-2 in the angiopoietin-tie signaling pathway. Cold Spring Harb Perspect Med 2, a006550 (2012).

37. Proescholdt MA, Jacobson S, Tresser N, Oldfield EH, Merrill MJ. Vascular Endothelial Growth Factor Is Expressed in Multiple Sclerosis Plaques and Can Induce Inflammatory Lesions in Experimental Allergic Encephalomyelitis Rats. Journal of Neuropathology & Experimental Neurology 61, 914–925 (2002).

38. Zhao X, Eyo UB, Murugan M, Wu L-J. Microglial interactions with the neurovascular system in physiology and pathology. Developmental neurobiology 78, 604–617 (2018).

39. Davalos D, et al. Fibrinogen-induced perivascular microglial clustering is required for the development of axonal damage in neuroinflammation. Nature communications 3, 1227 (2012).

40. Amstad A, et al. Growth differentiation factor 15 is increased in stable MS. Neurol Neuroimmunol Neuroinflamm 7, (2020).

41. Ivan DC, et al. Insulin-like growth factor-1 receptor controls the function of CNS-resident macrophages and their contribution to neuroinflammation. Acta neuropathologica communications 11, 35 (2023).

42. Jolly S, et al. G protein-coupled receptor 37-like 1 modulates astrocyte glutamate transporters and neuronal NMDA receptors and is neuroprotective in ischemia. Glia 66, 47–61 (2018).

43. Shao X, et al. Knowledge-graph-based cell-cell communication inference for spatially resolved transcriptomic data with SpaTalk. Nature communications 13, 4429 (2022).

44. Kaunzner UW, et al. Quantitative susceptibility mapping identifies inflammation in a subset of chronic multiple sclerosis lesions. Brain 142, 133–145 (2019).

45. Morabito S, et al. Single-nucleus chromatin accessibility and transcriptomic characterization of Alzheimer’s disease. Nat Genet 53, 1143–1155 (2021).

46. Morabito S, Reese F, Rahimzadeh N, Miyoshi E, Swarup V. hdWGCNA identifies co-expression networks in high-dimensional transcriptomics data. Cell Reports Methods 3, (2023).

47. Langfelder P, Luo R, Oldham MC, Horvath S. Is my network module preserved and reproducible? PLoS Comput Biol 7, e1001057 (2011).

48. Bayraktar OA, et al. Astrocyte layers in the mammalian cerebral cortex revealed by a single-cell in situ transcriptomic map. Nature Neuroscience, (2020).

49. Hendrickx DAE, et al. Gene Expression Profiling of Multiple Sclerosis Pathology Identifies Early Patterns of Demyelination Surrounding Chronic Active Lesions. Front Immunol 8, 1810 (2017).

50. Ponath G, Park C, Pitt D. The Role of Astrocytes in Multiple Sclerosis. Front Immunol 9, 217 (2018).

51. Wheeler MA, et al. MAFG-driven astrocytes promote CNS inflammation. Nature, (2020).

52. Wheeler MA, et al. Droplet-based forward genetic screening of astrocyte-microglia cross-talk. Science 379, 1023–1030 (2023).

53. McNamara NB, et al. Microglia regulate central nervous system myelin growth and integrity. Nature, (2022).

54. Webster HdF. Growth factors and myelin regeneration in multiple sclerosis. Multiple Sclerosis 3, 113–120 (1997).

55. Beenken A, Mohammadi M. The FGF family: biology, pathophysiology and therapy. Nat Rev Drug Discov 8, 235–253 (2009).

56. Clemente D, Ortega MC, Arenzana FJ, de Castro F. FGF-2 and Anosmin-1 are selectively expressed in different types of multiple sclerosis lesions. J Neurosci 31, 14899–14909 (2011).

57. Mohan H, et al. Transcript profiling of different types of multiple sclerosis lesions yields FGF1 as a promoter of remyelination. Acta neuropathologica communications 2, 168 (2014).

58. Furusho M, Roulois AJ, Franklin RJ, Bansal R. Fibroblast growth factor signaling in oligodendrocyte-lineage cells facilitates recovery of chronically demyelinated lesions but is redundant in acute lesions. Glia 63, 1714–1728 (2015).

59. Rajendran R, et al. The small molecule fibroblast growth factor receptor inhibitor infigratinib exerts anti-inflammatory effects and remyelination in a model of multiple sclerosis. Br J Pharmacol 180, 2989–3007 (2023).

60. Kotter MR, Li WW, Zhao C, Franklin RJ. Myelin impairs CNS remyelination by inhibiting oligodendrocyte precursor cell differentiation. J Neurosci 26, 328–332 (2006).

61. Schaffner E, et al. Myelin insulation as a risk factor for axonal degeneration in autoimmune demyelinating disease. Nat Neurosci 26, 1218–1228 (2023).

62. Pereira JB, et al. Microglial activation protects against accumulation of tau aggregates in nondemented individuals with underlying Alzheimer’s disease pathology. Nat Aging 2, 1138–1144 (2022).

63. Bellan M, Pirisi M, Sainaghi PP. The Gas6/TAM System and Multiple Sclerosis. Int J Mol Sci 17, (2016).

64. Weinger JG, Omari KM, Marsden K, Raine CS, Shafit-Zagardo B. Up-regulation of soluble Axl and Mer receptor tyrosine kinases negatively correlates with Gas6 in established multiple sclerosis lesions. The American journal of pathology 175, 283–293 (2009).

65. Cignarella F, et al. TREM2 activation on microglia promotes myelin debris clearance and remyelination in a model of multiple sclerosis. Acta Neuropathol, (2020).

66. Kaufmann M, et al. Identification of early neurodegenerative pathways in progressive multiple sclerosis. Nat Neurosci 25, 944–955 (2022).

67. Skripuletz T, Gudi V, Hackstette D, Stangel M. De- and remyelination in the CNS white and grey matter induced by cuprizone: the old, the new, and the unexpected. Histol Histopathol 26, 1585–1597 (2011).

68. Zhu J, et al. RNASequest: An End-to-End Reproducible RNAseq Data Analysis and Publishing Framework. Journal of Molecular Biology 435, 168017 (2023).

69. Hrvatin S, et al. A scalable platform for the development of cell-type-specific viral drivers. eLife 8, (2019).

70. Fleming SJ, et al. Unsupervised removal of systematic background noise from droplet-based single-cell experiments using CellBender. Nature Methods 20, 1323–1335 (2023).

71. Janssen P, et al. The effect of background noise and its removal on the analysis of single-cell expression data. Genome biology 24, 140 (2023).

72. Germain PL, Lun A, Garcia Meixide C, Macnair W, Robinson MD. Doublet identification in single-cell sequencing data using scDblFinder. F1000Res 10, 979 (2021).

73. Wolf FA, Angerer P, Theis FJ. SCANPY: large-scale single-cell gene expression data analysis. Genome biology 19, 15 (2018).

74. Bakken TE, et al. Single-nucleus and single-cell transcriptomes compared in matched cortical cell types. PLoS One 13, e0209648 (2018).

75. Hao Y, et al. Integrated analysis of multimodal single-cell data. Cell 184, 3573–3587.e3529 (2021).

76. Welch JD, Kozareva V, Ferreira A, Vanderburg C, Martin C, Macosko EZ. Single-Cell Multi-omic Integration Compares and Contrasts Features of Brain Cell Identity. Cell 177, 1873–1887.e1817 (2019).

77. Korsunsky I, et al. Fast, sensitive and accurate integration of single-cell data with Harmony. Nat Methods 16, 1289–1296 (2019).

78. Büttner M, Miao Z, Wolf FA, Teichmann SA, Theis FJ. A test metric for assessing single-cell RNA-seq batch correction. Nat Methods 16, 43–49 (2019).

79. He L, Davila-Velderrain J, Sumida TS, Hafler DA, Kellis M, Kulminski AM. NEBULA is a fast negative binomial mixed model for differential or co-expression analysis of large-scale multi-subject single-cell data. Commun Biol 4, 629 (2021).

80. Faure L, Soldatov R, Kharchenko PV, Adameyko I. scFates: a scalable python package for advanced pseudotime and bifurcation analysis from single-cell data. Bioinformatics 39, (2023).

81. Mao Q, Yang L, Wang L, Goodison S, Sun Y. SimplePPT: A Simple Principal Tree Algorithm.) (2015).

82. Albergante L, et al. Robust and Scalable Learning of Complex Intrinsic Dataset Geometry via ElPiGraph. Entropy (Basel) 22, (2020).

83. Zeira R, Land M, Strzalkowski A, Raphael BJ. Alignment and integration of spatial transcriptomics data. Nat Methods 19, 567–575 (2022).

84. Love MI, Huber W, Anders S. Moderated estimation of fold change and dispersion for RNA-seq data with DESeq2. Genome biology 15, 550 (2014).

85. Zhang Y, Parmigiani G, Johnson WE. ComBat-seq: batch effect adjustment for RNA-seq count data. NAR genomics and bioinformatics 2, lqaa078 (2020).

86. Zhang B, Horvath S. A general framework for weighted gene co-expression network analysis. Statistical applications in genetics and molecular biology 4, Article17 (2005).

