## Supplemental Table 1 for "Spatial transcriptomics reveals heterogeneous cell–cell interactions among brain regions in a cuprizone model consistent with multiple sclerosis lesions"

Table S1. Significant sender:receiver cell pairs and enriched receptor pathways in cortex predicted by SpaTalk

| **sender:receiver** | **receptor_pathways** |
| --- | --- |
| astrocytes:microglia | CRMPs in Sema3A signaling |
| astrocytes:microglia | Death Receptor Signalling |
| astrocytes:microglia | Herpes simplex virus 1 infection |
| astrocytes:microglia | Interleukin-12 family signaling |
| astrocytes:microglia | Interleukin-27 signaling |
| astrocytes:microglia | Late stage (branching morphogenesis) pancreatic bud precursor cells |
| astrocytes:microglia | MAP2K and MAPK activation |
| astrocytes:microglia | Pre-NOTCH Expression and Processing |
| astrocytes:microglia | Pre-NOTCH Processing in Golgi |
| astrocytes:microglia | Prion diseases |
| astrocytes:microglia | PTK6 promotes HIF1A stabilization |
| astrocytes:microglia | Regulation of Beta-Cell Development |
| astrocytes:microglia | Regulation of FZD by ubiquitination |
| astrocytes:microglia | Signaling by Non-Receptor Tyrosine Kinases |
| astrocytes:microglia | Signaling by NOTCH |
| astrocytes:microglia | Signaling by PTK6 |
| astrocytes:microglia | TNF signaling |
| astrocytes:microglia | TNFR1-induced NFkappaB signaling pathway |
| microglia:astrocytes | ECM-receptor interaction |
| microglia:mol | Disassembly of the destruction complex and recruitment of AXIN to the membrane |
| microglia:mol | Interleukin-12 family signaling |
| microglia:mol | Interleukin-27 signaling |
| microglia:mol | Interleukin-6 family signaling |
| microglia:mol | Interleukin-6 signaling |
| microglia:mol | Kaposi sarcoma-associated herpesvirus infection |
| microglia:mol | MAPK1 (ERK2) activation |
| microglia:mol | MAPK3 (ERK1) activation |
| microglia:mol | Parathyroid hormone synthesis, secretion and action |
| microglia:mol | RAF-independent MAPK1/3 activation |
| microglia:mol | TCF dependent signaling in response to WNT |
| microglia:mol | Wnt signaling pathway |
| microglia:opc | Growth hormone synthesis, secretion and action |
| microglia:opc | Human immunodeficiency virus 1 infection |
| microglia:opc | Interleukin-12 family signaling |
| microglia:opc | IRS activation |
| microglia:opc | Non-alcoholic fatty liver disease (NAFLD) |
| microglia:opc | PD-L1 expression and PD-1 checkpoint pathway in cancer |
| mol:microglia | Chemokine signaling pathway |
| mol:microglia | IRAK2 mediated activation of TAK1 complex upon TLR7/8 or 9 stimulation |
| mol:microglia | Regulation of FZD by ubiquitination |
| mol:microglia | TRAF6-mediated induction of TAK1 complex within TLR4 complex |
| mol:opc | Death Receptor Signalling |
| mol:opc | NOTCH3 Activation and Transmission of Signal to the Nucleus |
| mol:opc | p75 NTR receptor-mediated signalling |
| mol:opc | Signaling by NOTCH3 |
| opc:mol | Adherens junction |
| opc:mol | Cell adhesion molecules (CAMs) |
| opc:mol | ECM-receptor interaction |
| opc:mol | Signaling by MET |
