## Supplemental Table 2 for "Spatial transcriptomics reveals heterogeneous cell–cell interactions among brain regions in a cuprizone model consistent with multiple sclerosis lesions"

Table S2. Significant sender:receiver cell pairs and enriched receptor pathways in CC predicted by SpaTalk

| **snd_rec** | **receptor_pathways** |
| --- | --- |
| astrocytes:microglia | Endocrine resistance |
| astrocytes:microglia | Herpes simplex virus 1 infection |
| astrocytes:microglia | MAP2K and MAPK activation |
| astrocytes:microglia | Notch signaling pathway |
| astrocytes:microglia | Th1 and Th2 cell differentiation |
| microglia:astrocytes | Basal cell carcinoma |
| microglia:astrocytes | Melanogenesis |
| microglia:mol | Endocytosis |
| microglia:mol | JAK-STAT signaling pathway |
| microglia:opc | Bladder cancer |
| microglia:opc | Fibronectin matrix formation |
| microglia:opc | Signaling by Receptor Tyrosine Kinases |
| mol:microglia | Adipocytokine signaling pathway |
| mol:microglia | Amyotrophic lateral sclerosis (ALS) |
| mol:microglia | CRMPs in Sema3A signaling |
| mol:microglia | Endocrine resistance |
| mol:microglia | Noncanonical activation of NOTCH3 |
| mol:microglia | PTK6 promotes HIF1A stabilization |
| mol:microglia | Regulation of FZD by ubiquitination |
| mol:microglia | Signaling by Non-Receptor Tyrosine Kinases |
| mol:microglia | Signaling by PTK6 |
| mol:microglia | TNF signaling pathway |
| mol:microglia | TNFR2 non-canonical NF-kB pathway |
| mol:opc | Basal cell carcinoma |
| mol:opc | Cushing syndrome |
| mol:opc | RET signaling |
| opc:microglia | Endocrine resistance |
| opc:microglia | Fibronectin matrix formation |
| opc:microglia | Kaposi sarcoma-associated herpesvirus infection |
| opc:microglia | MAP2K and MAPK activation |
| opc:mol | Developmental Biology |
