## Supplementary figures and images for "Spatial transcriptomics reveals heterogeneous cell–cell interactions among brain regions in a cuprizone model consistent with multiple sclerosis lesions"

### Supplemental Figure 1

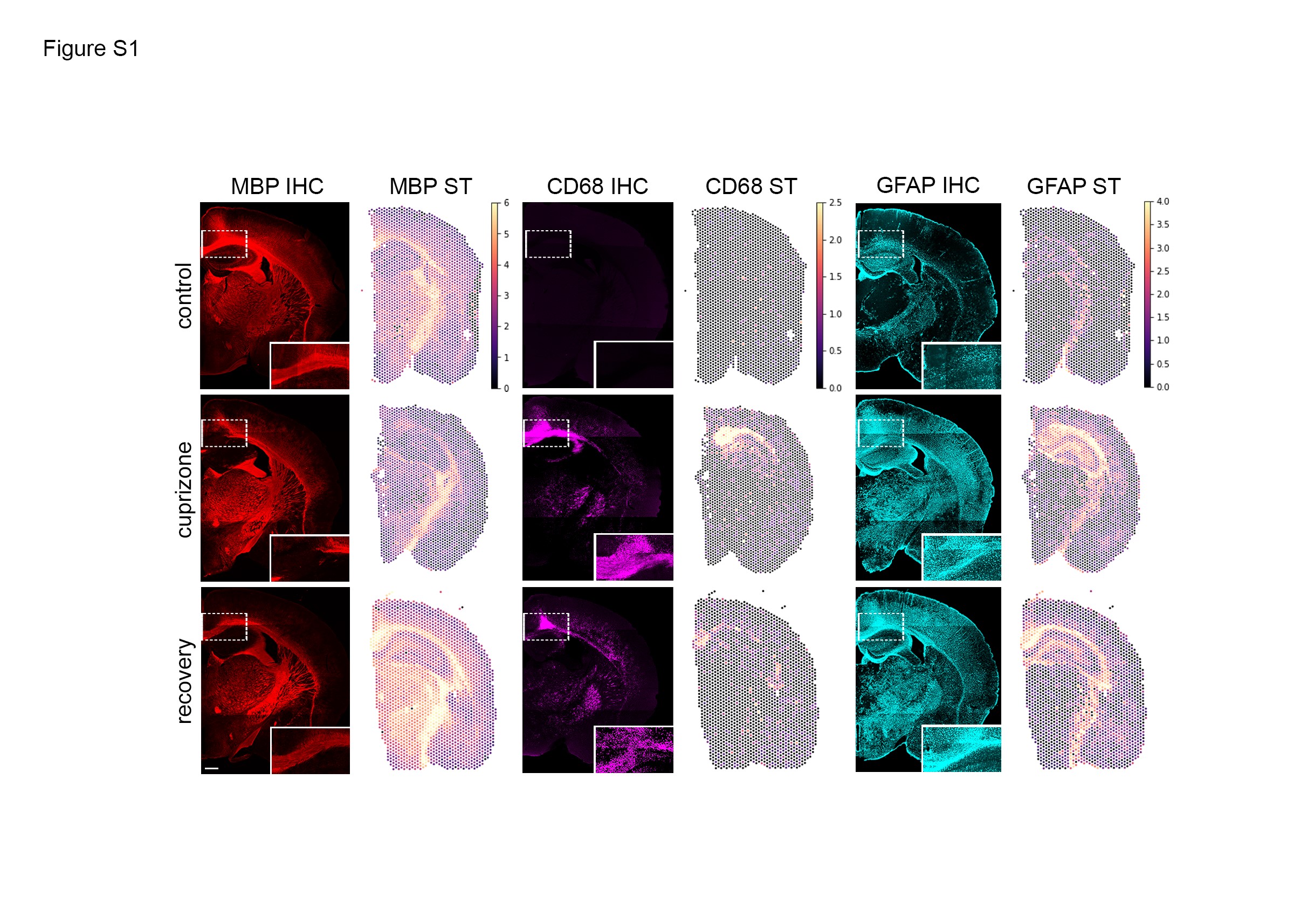

### Supplemental Figure 2

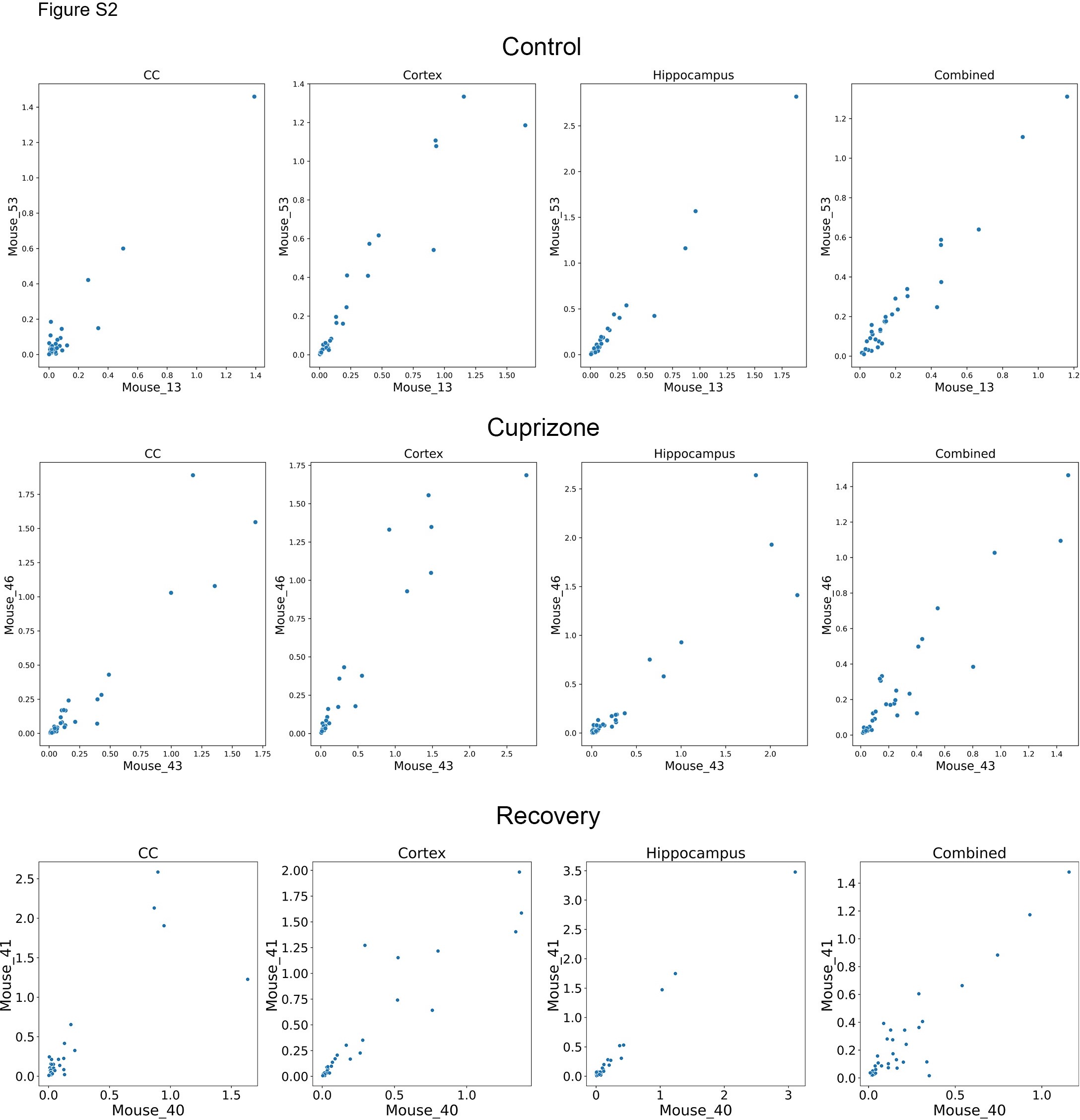

### Supplemental Figure 3

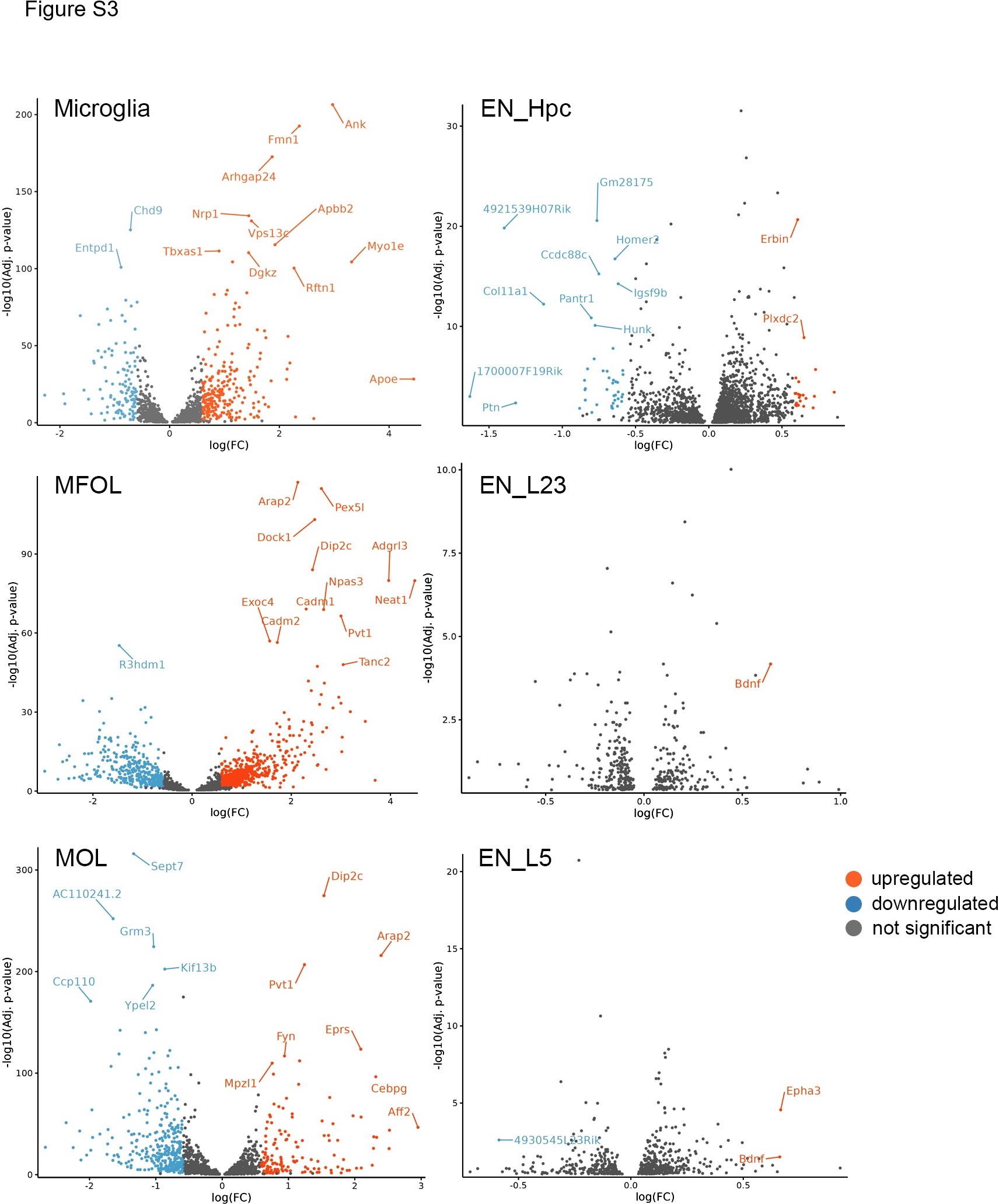

### Supplemental Figure 4

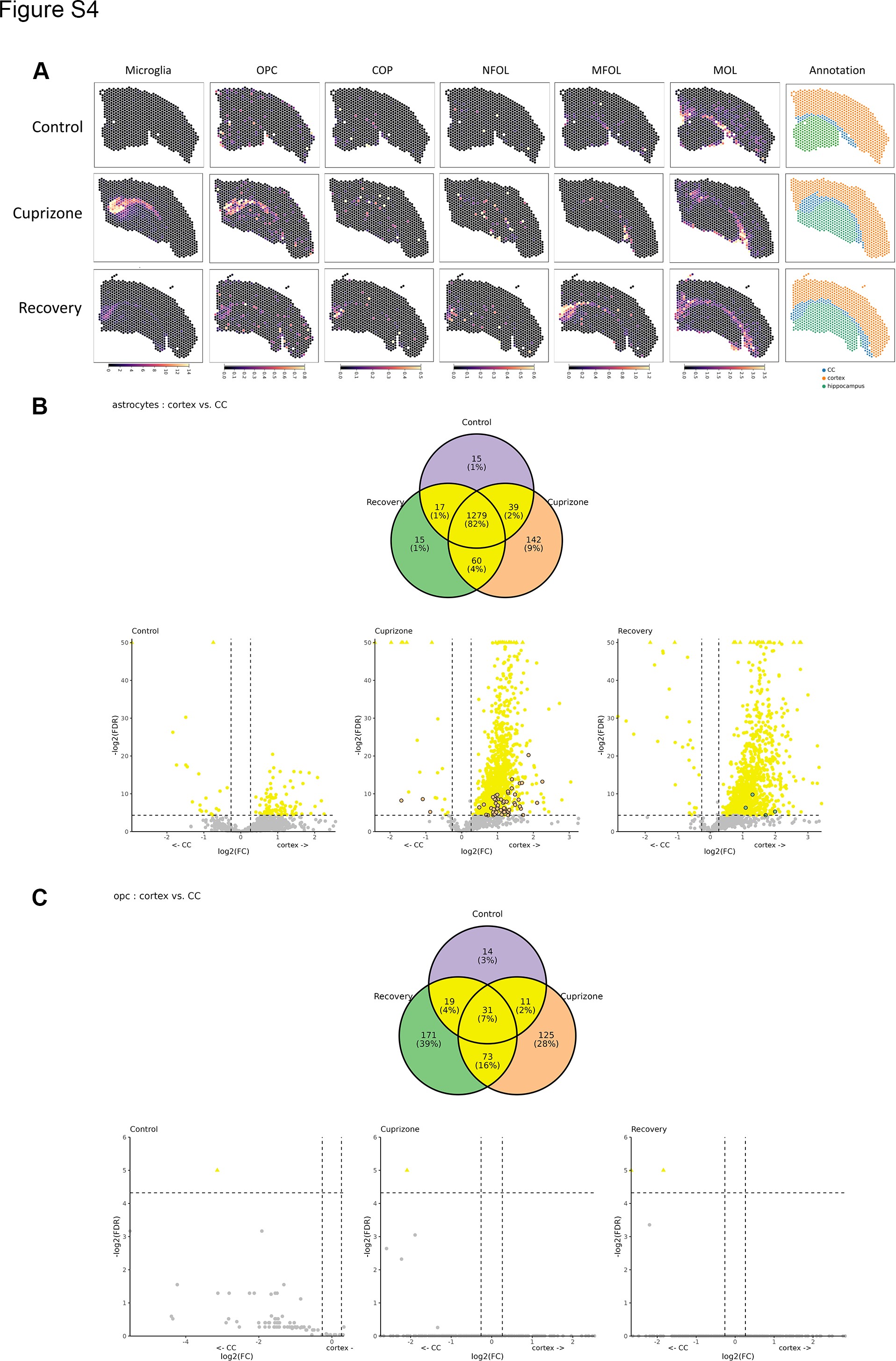

### Supplemental Figure 5

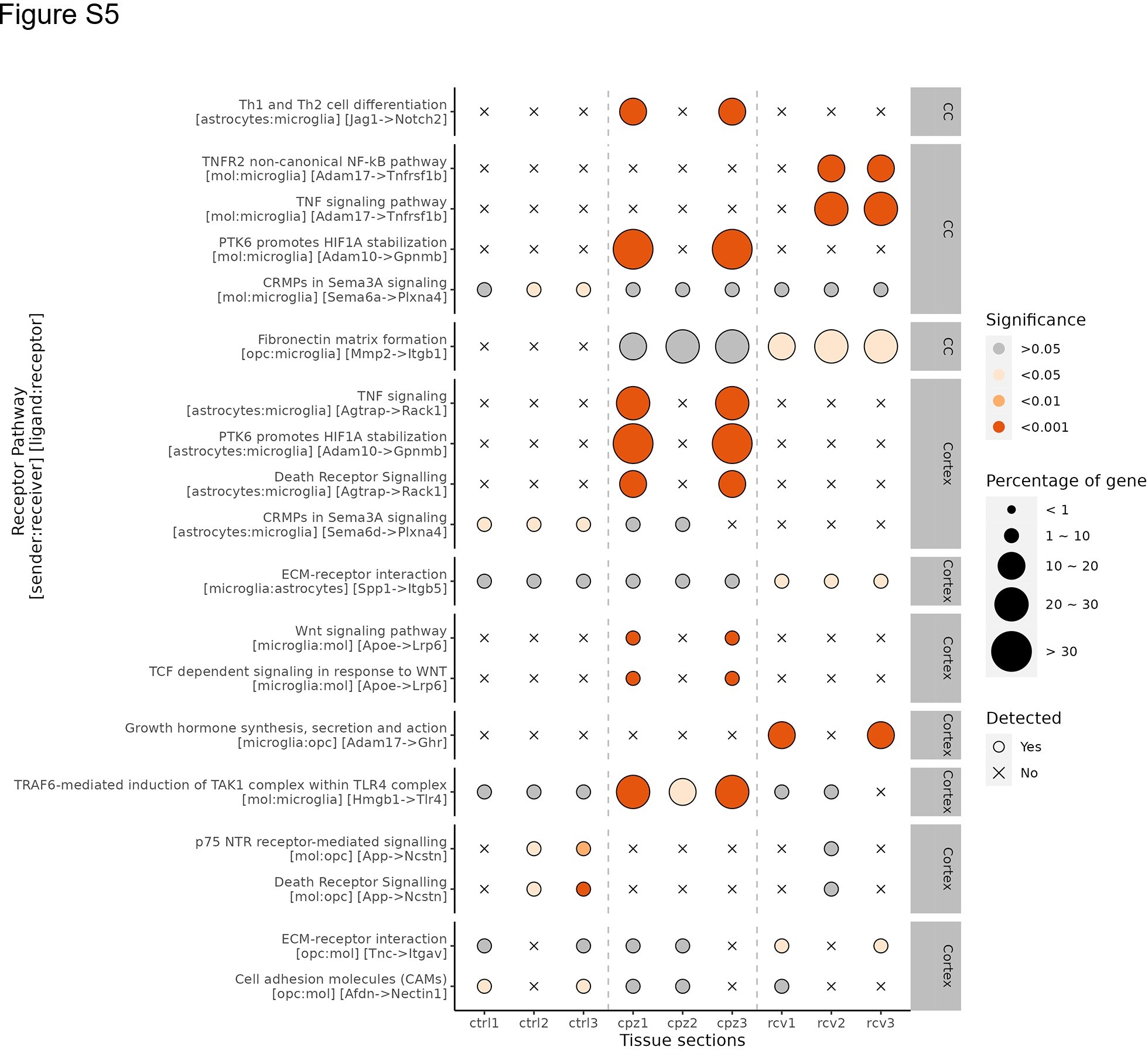
